# NETosis proceeds by cytoskeleton and endomembrane disassembly and PAD4-mediated chromatin de-condensation and nuclear envelope rupture

**DOI:** 10.1101/663427

**Authors:** Hawa Racine Thiam, Siu Ling Wong, Rong Qiu, Mark Kittisopikul, Amir Vahabikashi, Anne E. Goldman, Robert D. Goldman, Denisa D. Wagner, Clare M. Waterman

## Abstract

Neutrophil extracellular traps (NETs) are web-like DNA structures decorated with histones and cytotoxic proteins that are released by activated neutrophils to trap and neutralize pathogens during the innate immune response, but also form in and exacerbate sterile inflammation. Peptidylarginine deiminase 4 (PAD4) citrullinates histones and is required for NET formation (NETosis) in mouse neutrophils. While the *in vivo* impact of NETs is accumulating, the cellular events driving NETosis and the role of PAD4 in these events are unclear. We performed high resolution time-lapse microscopy of mouse and human neutrophils (PMN) and differentiated HL-60 neutrophil-like cells (dHL-60) labelled with fluorescent markers of organelles and stimulated with ionomycin or lipopolysaccharides to induce NETosis. Upon stimulation, cells exhibited rapid disassembly of the actin cytoskeleton, followed by shedding of plasma membrane microvesicles, disassembly and remodeling of the microtubule and vimentin cytoskeletons, ER vesiculation, chromatin de-condensation and nuclear rounding, progressive plasma membrane and nuclear envelope (NE) permeabilization, nuclear lamin meshwork and then NE rupture to release DNA into the cytoplasm, and finally plasma membrane rupture and discharge of extracellular DNA. Inhibition of actin disassembly blocked NET release. Mouse and dHL-60 cells bearing genetic alteration of PAD4 showed that chromatin de-condensation, lamin meshwork and NE rupture and extracellular DNA release required the enzymatic and nuclear localization activities of PAD4. Thus, NETosis proceeds by a step-wise sequence of cellular events culminating in the PAD4-mediated expulsion of DNA.

**Significance Statement:** Neutrophils are white blood cells specialized as the first line of host defense in the immune system. One way they protect organisms is through NETosis, in which they expel their DNA to form a web-like trap that ensnares pathogens and promotes clotting. However, NETs also mediate sterile inflammation, causing damage to the body. We used high-resolution live-cell microscopy to perform the first systematic characterization of the timing of dynamic cellular events leading to NETosis in human and mouse neutrophils and a neutrophil-like cell line. We discovered that NETosis proceeds by a step-wise sequence of cellular events that is conserved across species, and requires the activity of the PAD4 enzyme for DNA to be released from the nucleus and cell membrane.

## Introduction

Neutrophil granulocytes are central to innate immunity. Produced in the bone marrow at about a billion per kg of body mass (1), mature neutrophils are released to the circulation from where they are recruited to sites of injury by activated endothelial cells or resident macrophages (2). At injury sites, neutrophils deploy a variety of machineries to fight infection and neutralize pathogens including phagocytosis and degranulation, as well as the more recently characterized release of neutrophil extracellular traps (NETs) during NETosis (1, 2). NETs are web-like DNA structures decorated with histones and antimicrobial proteins that are released from stimulated neutrophils. NETs have been shown *in vitro* to trap and neutralize or kill pathogens including bacteria (3), fungi (4) and viruses (5) in a DNA scaffold and histone-dependent manner (3, 6). NETs have been demonstrated to limit bacterial growth *in vivo* in some model systems (6, 7) and to propagate inflammatory and immune responses (8, 9), positioning NETosis as an important part of innate immunity. However, soon after its discovery, NETosis was shown to convey detrimental effects, including endothelium and tissue damage during sepsis (10, 11) and mediation of thrombosis (12, 13). Furthermore, several autoimmune diseases are associated with high rates of NETosis and/or defects in NET clearance (11). More recently, NETs release has been proposed to promote cancer-associated thrombosis (14), tumorigenic microenvironments, and cancer cell migration (11, 15, 16) and to be involved in the awakening of dormant tumor cells in mouse models of prostate and breast cancer (17). Thus, understanding the cellular and molecular mechanisms mediating NETosis could facilitate either therapeutic improvement of innate immunity or mitigation of the damaging or disease-associated effects of NETosis.

The molecular requirements for NETosis have begun to be elucidated. NETosis can be stimulated in a variety of ways, including physiologically with various species of bacteria or yeast, monosodium urate crystals associated with gout, platelet activating factor, ionophores or bacterial lipopolysaccharides, or can be pharmacologically induced with phorbol ester (18). Depending on the stimuli, different downstream signaling pathways can mediate NETosis including PKC, calcium, Raf-MEK-ERK (19) that induces NADPH-MPO (20, 21) activation and ROS production but also the SYK-PI3K-mTorc2 pathway that mediates a ROS independent NETosis (22). Regardless of the stimulus, completion of NETosis requires convergence of signaling pathways on a mechanism that mediates the chromatin de-condensation that is necessary for NET release (23). Neutrophil elastase and other serine proteases contained in azurophilic granules have been proposed to mediate histone cleavage to facilitate their release from DNA (24, 25). PAD4, an enzyme that converts arginine to citrulline, is highly expressed in granulocytes (26, 27) as well as some cancer cells (28) and is the only member of the PAD family that possesses a nuclear localization signal (29). First described as a nuclear resident protein (29), PAD4 was recently shown to citrullinate cytoplasmic proteins (30) suggesting that it also localizes to the cytosol (26, 31, 32). During NETosis, PAD4 is proposed to translocate to the nucleus (11, 31) where it citrullinates histones, changing negatively charged arginines to neutral citrullines and thus reducing their charge-based interaction with DNA to promote chromatin de-condensation (23, 33). The relative importance of neutrophil elastase or PAD4 for completion of NETosis may be dictated by the cellular stimulus (24) or the species. Indeed, neutrophil elastase is required downstream of the NADPH pathway when NETosis is induced in human neutrophils by phorbol esters or Candida albicans (24, 25), while PAD4 is critical for NETosis in mouse neutrophils stimulated with calcium ionophore or certain strains of bacteria (33, 34). However, whether different cellular mechanisms are engaged during NETosis in mouse and human neutrophils and whether PAD4 is required for NET release in human neutrophils remains unclear.

In spite of the advancing knowledge of the molecular requirements of NETosis, less is known about the cellular mechanisms mediating this dramatic process (21). For DNA to be released to the cell exterior during NETosis, it must escape from the nucleus, pass through the cytoplasm containing an extensive network of membrane-bounded organelles and cytoskeletal systems and finally breach the plasma membrane. The expulsion of NETs is thought to result in neutrophil death (3, 21), but some other evidence suggests that neutrophils may survive NETosis and DNA-depleted cells could retain some phagocytic capacity and induce adaptive immune responses in what has been termed “vital NETosis” (6, 35, 36). While it is generally thought that decondensed chromatin is expelled via nuclear envelope and plasma membrane rupture (21), it has also been proposed that cytoplasmic vesicles containing DNA might be exocytosed to allow cell survival after NET release (6, 35). However, little is known about how chromatin breaches organelles and the cytoskeleton to pass through the cytoplasm. There is evidence that the actin cytoskeleton (37-39) and microtubules (38, 40) may disassemble during NEtosis, yet pharmacological perturbations show that either disassembly or stabilization of actin impairs NETosis (21, 37, 38), while microtubule disruption has little effect (38). Thus, how cytoskeletal systems dynamically remodel to allow passage of chromatin through the cytoplasm before extracellular release is not well understood. To date, most knowledge about NETosis was gleaned using fixed time point assays. However live cell imaging has proven to be important for understanding the dynamics and cellular complexity of NETosis (6, 18, 21, 22, 38). Indeed, it allowed the first analysis of the dynamic changes in chromatin compaction and plasma membrane composition and integrity during NETosis (18, 21, 22, 38). Thus, an in depth understanding of the cellular events driving NETosis requires the detailed visualization of various cellular compartments during the entire process.

Here we used time-lapse high resolution multi-mode microscopy of human and mouse blood-derived neutrophils (PMN) as well as differentiated human leukemia (dHL-60) neutrophil-like cells bearing fluorescent markers of organelle and cytoskeletal systems to quantitatively characterize the cellular events mediating NETosis after calcium ionophore or LPS stimulation. We demonstrate that NETosis proceeds by a well-defined and conserved sequence of cellular events. Characterization of neutrophils derived from PAD4 knockout mice or gene-edited dHL-60 cells demonstrates that PAD4 enzymatic activity and nuclear localization signal are required for efficient DNA de-condensation, nuclear lamin meshwork and nuclear envelope (NE) rupture and extracellular DNA release.

## RESULTS

### NETosis starts with plasma membrane microvesicle shedding prior to DNA de-condensation, DNA release into the cytoplasm, and extracellular DNA expulsion in mouse, human and differentiated HL60 neutrophil-like cells

To characterize the cellular events mediating neutrophil NETosis, we performed high-resolution time-lapse imaging of SiR-DNA-stained, blood-derived mouse and human polymorphonuclear neutrophils (PMN) isolated from fresh blood, as well as differentiated HL60 neutrophil-like cells (dHL60) during stimulation of NETosis with ionomycin. Cells were plated onto non-coated, gamma-irradiated coverslips for 5 minutes before imaging, and ionomycin (4 μM) was added 10 minutes after imaging commenced. We followed cell shape and membrane dynamics using differential interference contrast (DIC) microscopy, while spinning-disk confocal microscopy of the live-cell nuclear stain SiR-DNA allowed visualization of nuclear shape and DNA texture dynamics. Pairs of DIC and confocal images were acquired at two confocal planes at each 1-2 min time interval for 4 hr. One plane was at the ventral cell surface at the interface between the cell and coverslip and the second plane was at 3 μm above the coverslip in the cell center. Under these conditions, 15.5% of mouse PMNs, 44.6% of human PMNs and 55.6% of dHL60 cells observed went on to complete NETosis, as evidenced by extracellular expulsion of DNA (figure 1B, D and F). The NETosis efficiencies attained during live-cell imaging were similar to those measured in a fixed-timepoint assay at 2.5 hr after ionomycin stimulation (31% in dHL60; Supplementary figure 1), and those previously published for mouse (41) and human (24) PMNs, indicating that our long-term imaging is minimally perturbative to NETosis.

**Figure 1:**
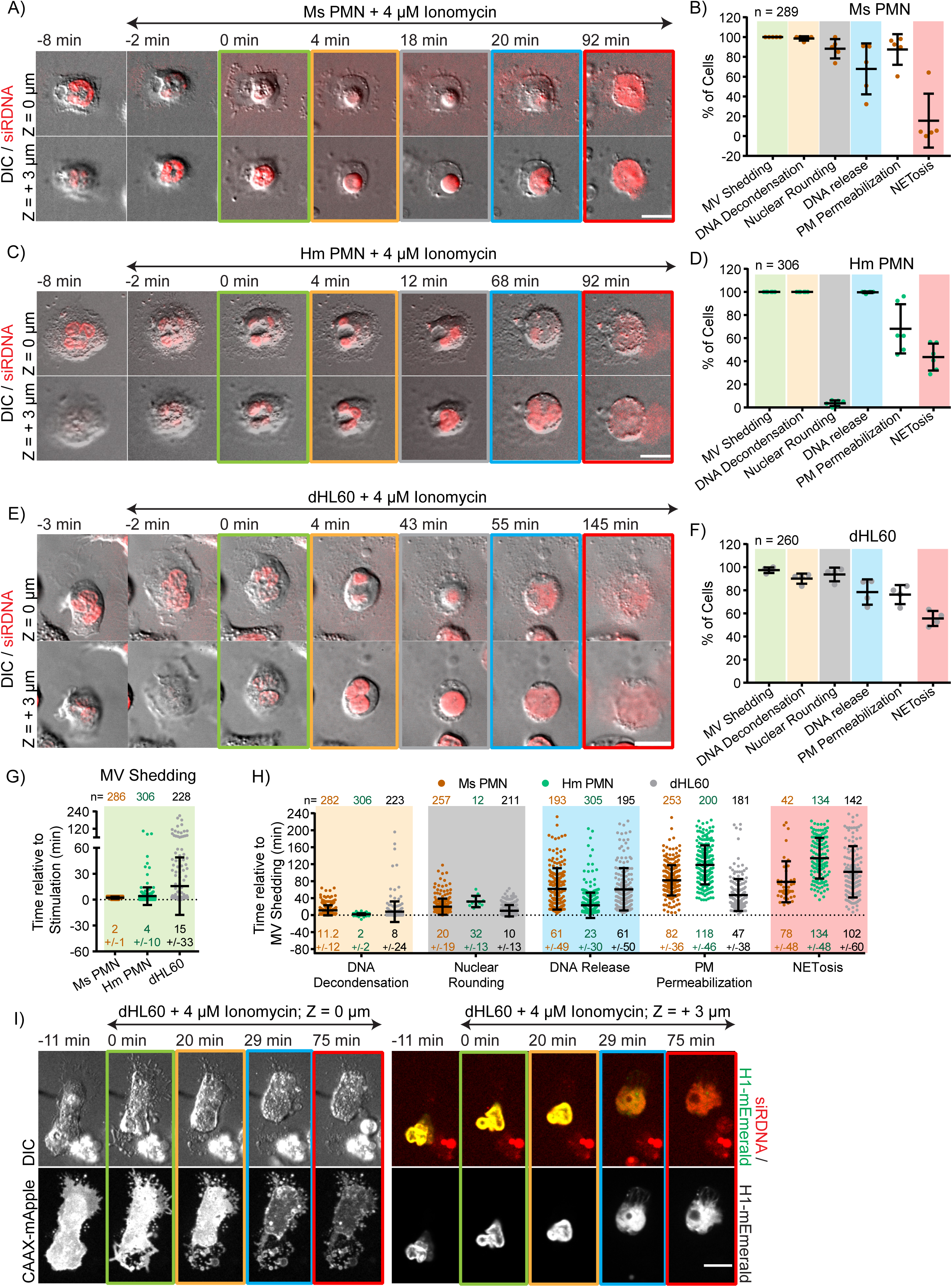
NETosis proceeds by plasma membrane microvesicle shedding, DNA de-condensation, DNA release from the nucleus into the cytoplasm, and extracellular DNA expulsion in mouse and human blood neutrophils and dHL60 cells. Mouse (Ms; A, B, G, H) or human (Hm; C, D, G, H) blood polymorphonuclear neutrophils (PMN) or differentiated HL60 neutrophil-like cells dHL60 cells (E, F, G, I) were stained with far-red SiR-DNA to fluorescently label DNA, plated on coverslips, stimulated with 4 μM ionomycin, and imaged live by differential interference contrast (DIC) and spinning-disk confocal microscopy at the coverslip-cell interface (Z = 0 μm) and 3 μm above in the cell center (Z = +3 μm) at 1-2 min intervals for 4 hours, with time in min relative to plasma membrane microvesicle shedding noted above images. Double-arrow-headed lines above images indicate cells that are in presence of ionomycin, boxes around images correspond to cellular event categories in the graphs, with the color of the box indicating the event with the same colored background in graphs. A, C, E: Time series of image overlays of DIC (greyscale) and fluorescence (red) of cell and DNA dynamics. B, D, F: Quantification of the percent of cells that exhibit microvesicle (MV) shedding, DNA de-condensation, nuclear rounding, release of nuclear DNA into the cytoplasm (DNA Release), loss of DIC contrast in the cell periphery suggesting plasma membrane permeabilization (PM Permeabilization), and extracellular DNA release (NETosis) after ionomycin stimulation determined from time-lapse DIC and confocal color-overlay movies. n = total number of cell observed, each point represents the percent of cells performing the event in one experiment, long bar = mean, short bars = S.D. G, H: Quantification of the timing of microvesicle shedding relative to ionomycin stimulation (G) or of cellular event initiation relative to microvesicle shedding (H) determined from time-lapse DIC and confocal color-overlay movies. n= total number of cells observed, points represent individual cells, mean (long bar) and S.D. (short bar) also shown below each plot. I: Time series of DIC (upper row-left) and fluorescence images of cell, plasma membrane (lower row-left) and histone H1 (right: upper row, green; lower row) dynamics in a living dHL60 cell stained with far-red SiR-DNA and co-expressing the farnesylation signal sequence CAAX fused to mApple (CAAX-mApple) as a plasma membrane marker and H1-mEmerald as a histone marker. Boxes around images indicate different cellular events (green: MV Shedding; yellow: DNA De-condensation; blue: DNA release to the cytosol, red: extracellular DNA release). A, C, E, I: bars = 10 μm.

We then examined time-lapse movies to analyze cell dynamics during NETosis. DIC movies of the cell-coverslip interface of all three cell types revealed that within minutes after ionomycin stimulation, cells rapidly formed and shed many small (∼< 2 μm) membrane vesicles (figure 1A, C, E, G, and i; Movie 1). The plasma membrane origin of the microvesicles was confirmed by imaging ionomycin-stimulated dHL60 cells expressing the farnesylation signal sequence CAAX fused to mApple (CAAX-mApple) as a plasma membrane marker (figure 1I, Movie 2), and similar plasma membrane vesiculation was observed in mouse PMNs stimulated with lipopolysaccharide (LPS, 25 μg/mL) (Supplementary figure 2A, Movie 1). Following such microvesicle shedding, cells rounded up and eventually expelled extracellular DNA marking completion of NETosis. Quantification revealed that although not all cells that shed plasma membrane microvesicles went on to expel DNA, 100% of cells that completed NETosis had previously formed and shed microvesicles. Thus, plasma membrane microvesicle shedding is a hallmark of cellular entry in to the process of NETosis (figure 1B, D, F and I), and shall be used as an easily visualized temporal initiation point of NETosis for the remainder of this study.

We next compared nuclear and membrane dynamics by examining SiR-DNA and DIC movies in the cells’ central plane. Prior to ionomycin stimulation, nuclei of all three cell types were highly lobulated. The fluorescent DNA staining within cell nuclei was spatially heterogeneous with brighter and dimmer regions, indicative of regions of varied levels of chromatin condensation. Quantification of timing in movies showed that within ∼10min after ionomycin-induced plasma membrane microvesicle shedding, the spatial heterogeneity of DNA fluorescence reduced and became homogeneous throughout the nucleus (figure 1H, Movie 1). Similar homogenization of DNA fluorescence was observed during NETosis in dHL60 cells expressing mEmerald-tagged linker histone H1 (histone-mEmerald, figure 1I, Movie 2), suggesting that these changes in the distribution of fluorescence correspond to chromatin de-condensation. De-condensation was followed ∼10min later by loss of nuclear lobulation and the rounding of the nuclear surface in both mouse PMN and dHL60, while in human PMN, nuclei de-lobulated and partially rounded (figure 1C, H). Approximately one hour (for mouse PMN and dHL60) or 20min (for human PMN) after microvesicle shedding, the sharp DIC contrast band at the rim of the nucleus was lost and DNA expanded rapidly from the nucleus into the cytosol, suggesting nuclear rupture (figure 1H, Movie 1). There was also a sudden reduction in DIC contrast in the cell periphery (Movie 3) indicating a decrease in refractive index that likely corresponded to at least partial loss of cellular contents prior to nuclear rupture. In spite of this apparent leakage of cytosol, the peripheral cell membrane appeared intact at the resolution of the light microscope, and DNA was not released extracellularly until many minutes later, marking the completion of NETosis (figure 1B, D, F, H and I). Although not all stimulated cells that exhibited DNA de-condensation and leakage of cytosol expelled extracellular DNA to complete NETosis, 100% of cells that completed NETosis had previously completed these cellular processes. However, while all cells that completed NETosis released decondensed DNA to the cytoplasm prior to extracellular DNA release, a substantial portion of cells that released DNA into the cytoplasm did not go on to release it extracellularly (figure 1B, D, F). This suggests that DNA release into the cytoplasm may be required but not sufficient for extracellular DNA release. Together, these data indicate that NETosis proceeds by a dynamic, step-wise sequence of cellular events.

### ER vesiculation is followed by nuclear lamin meshwork disassembly, NE rupture and release of DNA into the cytoplasm leading to NETosis

For DNA to be released to the cell exterior during NETosis, it must escape from the nucleus, pass through the cytoplasm containing the cytoskeleton, and finally breach the plasma membrane. We initially focused on how DNA is released from the nucleus, which requires passing first through the meshwork of nuclear lamin isoforms located in the nuclear lamina, and then through both membranes of the nuclear envelope (NE). To examine lamin meshwork organization and dynamics, we first confirmed the expression and peripheral nuclear localization of lamin B1, B2 and A/C in dHL60 (42, 43) and of lamin B1 in human and mouse PMN by western blot and/or immunostaining (figure 2A,C, Supplementary figure 3A, B, C and D). Fixation and immunostaining of lamin B1/B2 in mouse PMN and lamin B1 in human PMN at time points after stimulation of NETosis with ionomycin showed discontinuities in the lamin meshwork detectable as early as 5 min after stimulation, with extrusion of DNA from the nucleus through the discontinuities and into the cytoplasm visible at 10 min, and disassembly of the lamin meshwork at later timepoints (figure 2A, B and C).

**Figure 2:**
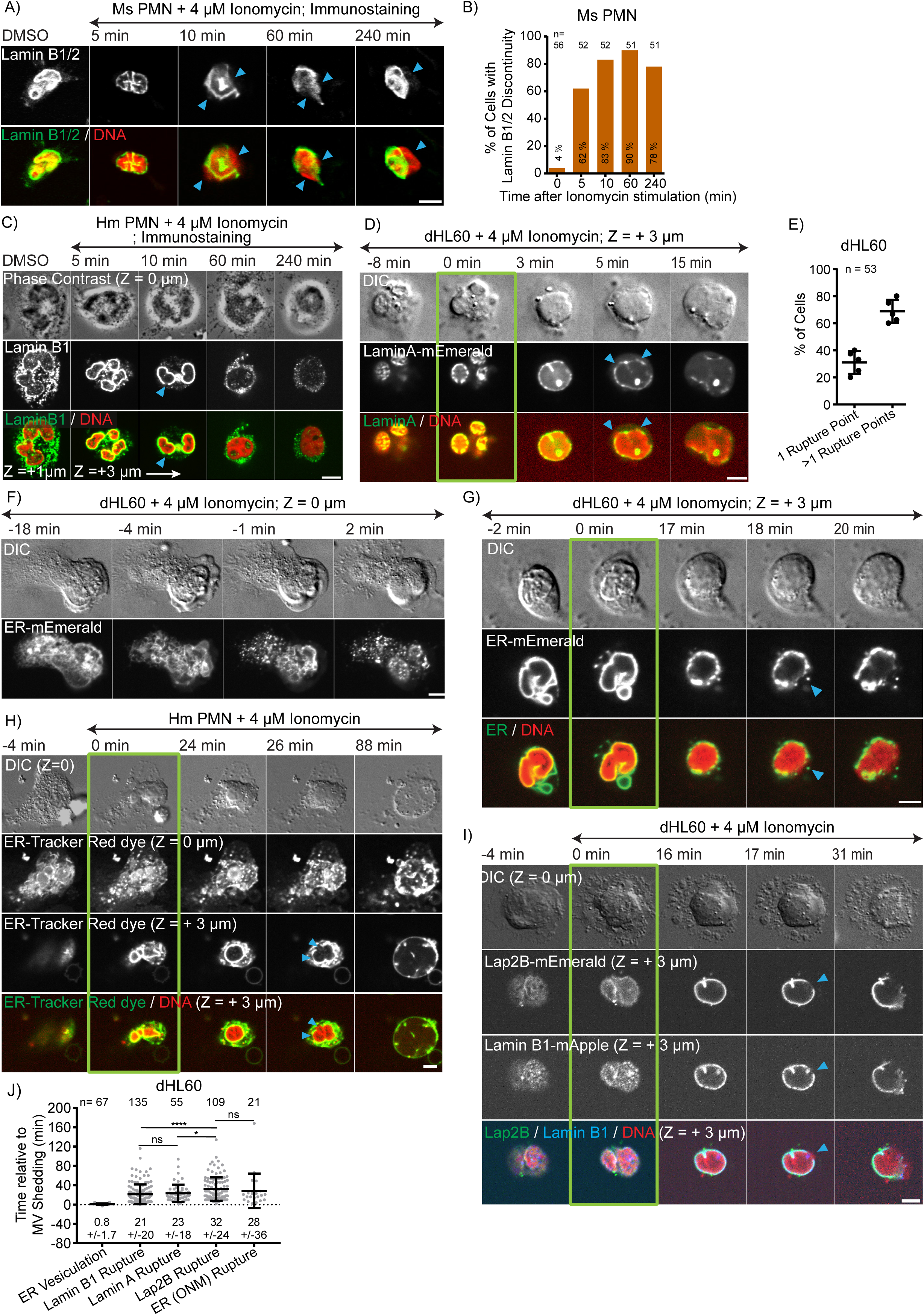
ER vesiculation is followed by nuclear lamina disassembly, NE rupture and release of DNA into the cytoplasm, leading to NETosis. Mouse (Ms; A, B) or human (Hm; C, H) blood polymorphonuclear neutrophils (PMN) or dHL60 cells (D, E, F, G, I, J) were plated on coverslips, stimulated with 4 μM ionomycin and fixed (panels A, C) at different time points after stimulation (noted on images) or imaged live (panels F,G,H, I) by differential interference contrast (DIC) and spinning-disk confocal microscopy at the coverslip-cell interface (Z = 0 μm) and 3 μm above in the cell center (Z = + 3 μm) at 1-2 min intervals for 4 hours, with time in min relative to plasma membrane microvesicle shedding noted above images. Double-arrow-headed lines above images indicate cells that are in presence of ionomycin, blue arrowheads mark rupture points in the lamina or NE. D, G-I: Green boxes around images indicate MV shedding. A: Immunofluorescence of lamin B1/B2 (upper row and bottom row, green) and DAPI (at 10 min) or SiR-DNA (DMSO, 5, 60 and 240 min) staining of DNA (bottom row, red) in fixed Ms PMN. B: Quantification from immunostaining experiments of the percent (numbers in bar) of Ms PMN exhibiting discontinuities in the lamin B1/B2 meshwork at different time points after stimulation, 0 represent cells treated with DMSO only. n = total number of observed cells. C: Immunofluorescence of lamin B1 (middle row and bottom row, green) and DAPI staining of DNA (bottom row, red) in Hm PMN fixed after 5 min of DMSO or 5, 10, 60 and 240 min of ionomyocin treatment. Z-stacks of 0.2 μm steps were acquired on a spinning-disk confocal microscope. Top row: Phase contrast images at the bottom plane (Z = 0 μm); Middle and bottom rows: Confocal images at the bottom plane (Z = 0 μm) and 1 or 3 μm above in the cell center (Z = + 1 μm, Z = + 3 μm). D: Time-series of DIC (top row) and confocal (lower rows) images of live dHL60 cells expressing lamin A-mEmerald (middle row and bottom row, green) and stained with far-red SiR-DNA to fluorescently label DNA (SiR-DNA, bottom row, red). E: Quantification of the percent of cells exhibiting rupture of the nuclear lamina at one or more than one points determined from movies of dHL60 cells expressing lamin A-mEmerald. n = total number of cell observed, each point represents the percent of cells in one experiment, long bar = mean, short bars = S.D. C: F, G, H: Time-series of DIC (upper rows) and confocal (lower rows) images of live dHL60 cells (F, G) expressing the KDEL sequence combined with the ER-retention signal sequence from calreticulin fused to mEmerald (ER-mEmerald) (F, bottom row; G, middle and bottom row, green) or live Hm PMN (H) stained with ER-Tracker Red (H, middle rows and bottom row, green) to label the ER and nuclear envelope (NE) and stained with SiR-DNA (G, H, bottom rows red). I: Time-series of DIC (upper rows) and confocal (lower rows) images of live dHL60 cells co-expressing the inner nuclear envelope protein, Lap2β-mEmerald (second row and bottom row, green) and lamin B1-mApple (third row and bottom row, blue) and stained with SiR-DNA (bottom row, red). In (G, H,I) the blue arrowheads highlight site of lamina and NE rupture. J: Quantification of the timing of the initiation of ER vesiculation or rupture of the nuclear lamina (Lamin B1 or Lamin A Rupture), the inner NE (Lap2β Rupture) or outer NE (ER (ONM) Rupture) relative to microvesicle shedding determined from time-lapse DIC and confocal color-overlay movies of dHL60 cells. n = total number of cell observed, points represent individual cells, mean (long bar) and S.D. (short bar) shown below each plot. **** and * for P value respectively < 0.0001, and <0.1; ns for non-significant. Statistical test: Mann-Whitney Test. A, C, D, F-I: bars = 5 μm.

To characterize the apparent difference in timing between the formation of lamin meshwork discontinuities and DNA release into the cytoplasm seen in fixed timepoint analysis of PMNs, we performed time-lapse DIC and spinning disk confocal microscopy on SiR-DNA-stained live dHL60 cells expressing either lamin A- or B1-mEmerald (Movie 4). Prior to stimulation, mEmerald-tagged lamins formed meshworks at the nuclear surface as seen in grazing optical sections near the ventral cell surface, and a somewhat punctate rim around the nuclear periphery in central optical sections (figure 2D, I, Supplementary figure 3B, C). Examination of movies after ionomycin stimulation showed that after microvesicle shedding, DNA de-condensation and nuclear rounding, the lamin meshwork acquired discontinuities often at multiple points (figure 2D, E and I). This was followed within 1-2 min by DNA extrusion through the lamin meshwork discontinuity into the cytosol. Initiation of the first detected discontinuity was soon followed by disassembly of the lamin meshwork as the DNA rapidly expanded throughout the cytoplasm (figure 2D, I, Movie 4). The apparent time lag between discontinuities in the lamin meshwork and release of nuclear DNA into the cytoplasm suggests that loss of lamin meshwork structural integrity may precede NE rupture.

We next analyzed NE and lamin dynamics simultaneously to determine the timing of the appearance of lamin meshwork discontinuities relative to NE rupture during NETosis. To this end, we exploited the continuity between the endoplasmic reticulum (ER) and NE to visualize the outer nuclear membrane by staining mouse and human PMN with a fluorescent ER vital dye (ER-dye, Movie 5) or by transiently expressing the KDEL sequence combined with the ER-retention signal sequence from calreticulin fused to mEmerald (ER-5-mEmerald) in dHL60 (Movie 6). We also expressed the inner nuclear envelope protein, Lap2β-mEmerald, in dHL60 (Movie 6). Time-lapse microscopy showed that prior to ionomycin stimulation, ER markers localized to a reticular network in the cell periphery as well as the NE, while Lap2β-mEmerald localized only to the NE. Upon ionomycin stimulation and soon after plasma membrane microvesicle shedding, the ER network rapidly broke into vesicles (figure 2F, G, H, J, Supplementary figure 4A, Movie 5) while the NE continued to stay intact around the nuclear periphery during rounding and chromatin de-condensation (figure 2G, H and supplementary figure 4A). Subsequent to nuclear rounding, the NE suddenly became discontinuous and rapidly retracted from the exact sites where DNA was extending from the nucleus into the cytoplasm, signifying NE rupture (figure 2G, H and supplementary figure 4A, Movie 6).

Co-expression of Lap2β-mEmerald and lamin B1-mApple in dHL60 showed that both the NE and lamin meshwork ruptured at the same location (Movie 7), while quantification of timing showed that the lamin meshwork became discontinuous slightly before NE rupture (figure 2I, J). Following such nuclear rupture, no NE or lamin meshwork re-sealing was ever observed. In addition, we never observed DNA to be released from the nucleus into the cytoplasm or extracellular space either alone or in NE-derived membrane vesicles (35) without prior lamin meshwork and NE rupture. Mouse PMN stimulated with LPS also vesiculated their ER after microvesicle shedding and prior to NE rupture and DNA release into the cytoplasm (Supplementary figure 2B, Movie 5). These results show that during NETosis, cells vesiculate their ER network before decondensing their DNA and rupturing their lamin meshwork at one or multiple points, followed by NE rupture to allow release of decondensed DNA into the cytoplasm. Furthermore, the distinct timings of ER vesiculation, lamin meshwork disassembly, and NE membrane rupture suggest they proceed under discrete regulatory control.

### The actin, microtubule and peripheral vimentin cytoskeletons disassemble prior to NETosis, and F-actin disassembly is important for NET release

We next sought to examine how cytoplasmic DNA breaches the actin, microtubule (MT) and vimentin intermediate filament (IF) cytoskeletal networks to eventually be expelled into the extracellular environment during NETosis. To visualize actin dynamics, we stained mouse or human PMN with the membrane-permeable actin filament marker SiR-actin (Movies 9 and 10) or stably expressed either F-tractin-mApple (Movie 8) or actin-mEmerald (Movie 11) to mark actin filaments or total actin (filaments and monomers), respectively (44) in dHL60. Examination of time-lapse movies revealed that prior to ionomycin stimulation, actin markers concentrated in punctate structures at the cell’s ventral surface as well as dynamic ruffling protrusions or contracting uropods at the periphery of spread neutrophils. Ionomycin stimulation induced the rapid dissociation of fluorescent actin markers from these structures and their redistribution to diffuse fluorescence throughout the cytosol, indicating filament disassembly (Movie 8, 9, 10). Comparison with the timing of microvesicle shedding showed that actin disassembly was completed on average ∼2 min prior to microvesicle shedding. Examination of individual cells showed that actin filament disassembly always closely preceded vesicle shedding, and was followed many minutes later by changes in DNA condensation and nuclear architecture (figure 3A, B, G, supplementary figure 4B). Similar disassembly of actin prior to microvesicle shedding was observed in SiR-actin-stained mouse PMN stimulated with LPS (supplementary figure 2C, Movie 10). Thus, actin cytoskeletal disassembly is the earliest cellular event in the process of NETosis.

MT dynamics were then analyzed during NETosis in ionomycin-stimulated mouse and human PMN stained with SiR-tubulin (Movie 9, 10) or dHL60 transiently expressing the ensconsin microtubule-binding domain fused to eGFP (eMTB-GFP, (45), Movie 8) or tubulin fused to mEmerald (Movie 12). Prior to stimulation, MTs were organized in a radial array with their (presumably) minus ends anchored at a perinuclear microtubule organizing center (MTOC) with their free (plus) ends located at the cell periphery. Time-lapse microscopy showed that soon after ionomycin stimulation the MT network rapidly disassembled from the free plus ends inwards to the MTOC where MT markers remained visible (Movies 8, 9, 10). Analysis and quantification indicated that MT disassembly was complete just after microvesicle shedding (figure 3C, D, G, supplementary figure 4C), but prior to DNA de-condensation or nuclear rounding. We confirmed that MT disassembly occurred with similar timing in mouse PMN undergoing NETosis after LPS stimulation (supplementary figure 2D, Movie 10).

**Figure 3:**
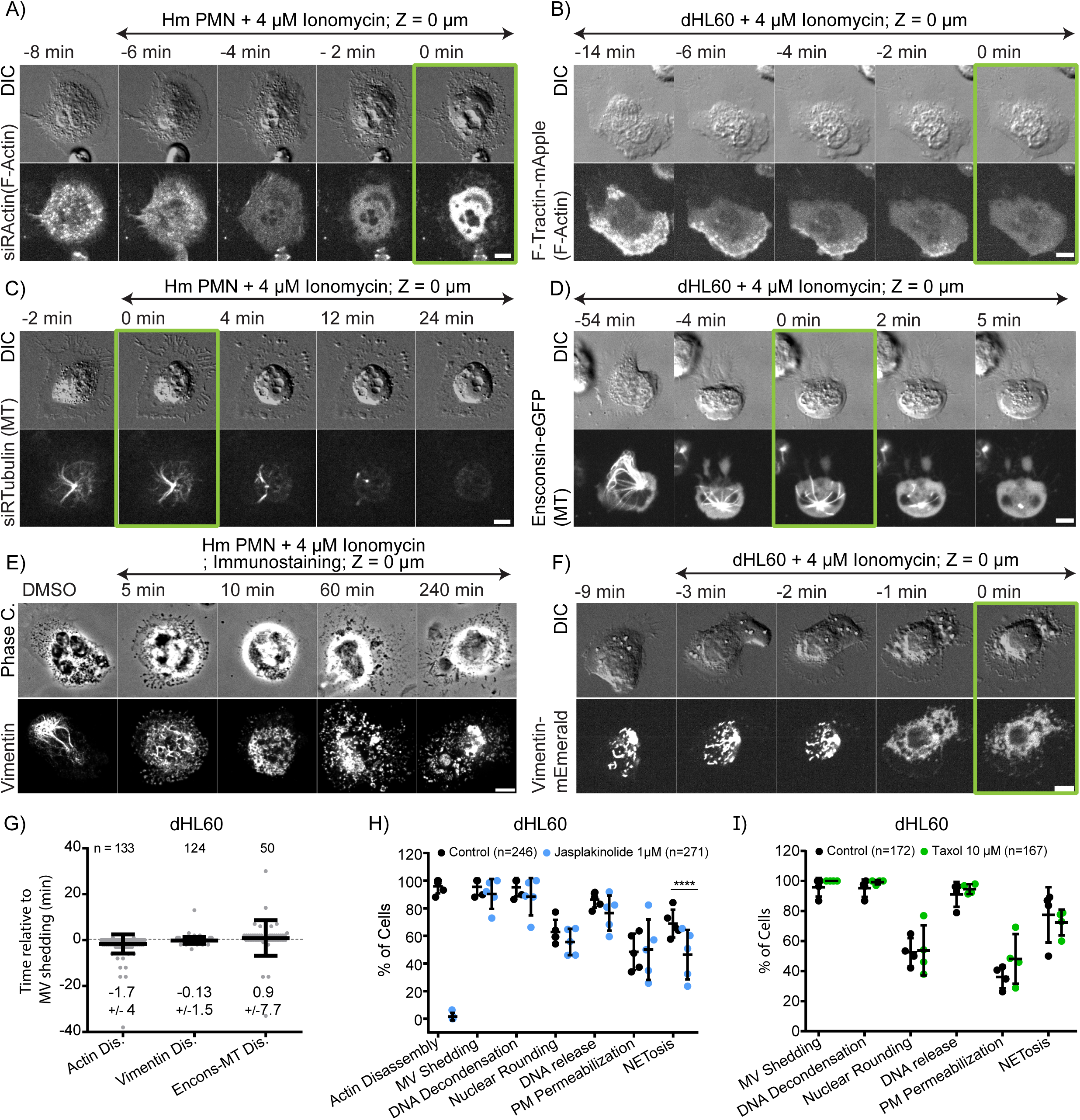
The actin, microtubule and vimentin cytoskeletons disassemble prior to NETosis, and F-actin disassembly is important for NET release. Human blood polymorphonuclear neutrophils (Hm PMN; A, C, E) or differentiated HL60 neutrophil-like cells (dHL60; B, D, F, G-I) were plated on coverslips, stimulated with 4 μM ionomycin and fixed at different time points after stimulation (noted above images, E) or imaged live by differential interference contrast (DIC) and spinning-disk confocal microscopy at the coverslip-cell interface (Z = 0 μm) and 3 μm above in the cell center (Z = +3 μm) at 1-2 min intervals for 4 hours, with time in min relative to plasma membrane microvesicle shedding noted above images. Double-arrow-headed lines above images indicate cells that are in presence of ionomycin. A-F: Time-series of DIC (A-D and F) or phase contrast (E) (top row) and confocal (lower rows) images of live (A, C) or fixed (E) Hm PMN or live dHL60 cells (B, D, F). A, B: Cells were stained with far red SiR-actin (A) or expressing F-Tractin-mApple (B) to fluorescently label actin filaments. C, D: Cells were stained with far red SiR-tubulin (C) or expressing the ensconsin microtubule binding domain fused to eGFP (D) to fluorescently label microtubules. E, F: Cells were fixed and vimentin was immunolocalized (E) or were expressing vimentin-mEmerald (F) to fluorescently label vimentin intermediate filaments. A-D, F: Green boxes around images indicate MV Shedding. G: Quantification of the timing of the initiation of actin and vimentin disassembly (Dis) and end of microtubule disassembly relative to microvesicle shedding determined from time-lapse DIC and confocal color-overlay movies of dHL60 cells. n = total number of cell observed, points represent individual cells, mean (long bar) and S.D. (short bar) shown below each plot. H, I: Quantification of the percent of cells that exhibit actin depolymerization (Actin Disassembly), microvesicle (MV) shedding, DNA de-condensation, nuclear rounding, release of nuclear DNA into the cytoplasm (DNA Release), loss of DIC contrast in the cell periphery suggesting plasma membrane permeabilization (PM Permeabilization), and extracellular DNA release (NETosis) after ionomycin stimulation determined from time-lapse DIC and confocal color-overlay movies of cells treated with vehicle (Control) or jasplakinolide (1 μM, H) or taxol (10 μM, I) to prevent actin and microtubule disassembly, respectively. n = total number of cell observed, points represent the percent of cells in one experiment, long bar = mean, short bars = S.D. Statistical test: Fisher’s Exact Test. A-F: bars = 5 μm.

The IF cytoskeleton was analyzed during NETosis in dHL60 cells transiently expressing vimentin-mEmerald (Movie 8). Prior to ionomycin stimulation, vimentin IFs were localized around the nuclear surface and there were filamentous and particulate vimentin structures in the cell periphery. After stimulation, peripheral vimentin-mEmerald structures rapidly disappeared and the fluorescence became diffuse, masking the ability to see perinuclear vimentin IF. Loss of peripheral vimentin structures was nearly concomitant with microvesicle shedding but before DNA de-condensation, nuclear rounding, or DNA release into the cytoplasm or extracellular expulsion (figure 3F and G, Movie 8). Vimentin IF organization was examined during NETosis in human and mouse PMN by fixation and immunostaining at different time points after ionomycin stimulation. This showed that peripheral vimentin IFs were largely disassembled within 5 min after stimulation and were absent from most observed cells 10 min after stimulation, while some dense and particulate vimentin staining remained near the cell center (figure 3E, supplementary figure 4I and J). Together these data showed that cells undergoing NETosis first disassemble their actin cytoskeleton, followed by microvesicle shedding, disassembly of microtubules and vimentin IFs prior to DNA de-condensation, nuclear rounding, nuclear lamin meshwork and NE rupture, DNA release into the cytoplasm, and extracellular DNA expulsion.

We then sought to determine the requirement for cytoskeletal disassembly in NETosis. To prevent disassembly, we used jasplakinolide or taxol, to stabilize F-actin or microtubules, respectively, and analyzed their effects on cytoskeleton organization and cells’ progression through NETosis. We did not test the role of vimentin IF disassembly in NETosis as there are no known methods of achieving vimentin IF stabilization. dHL60 cells expressing actin- or tubulin-mEmerald were plated for 10 min then pre-treated with 1 µM jasplakinolike or 10 µM taxol for 1 hr prior to ionomycin stimulation of NETosis. Inspection of the cytoskeletons showed that jasplakinolide treatment caused formation of a dense cortical actin network that remained mostly intact throughout NETosis (Movie 11), while taxol treatment blocked the disassembly of MTs (Movie 12) and induced peripheral MT bundles. Quantification showed that compared to vehicle control, neither jasplakinolide nor taxol had any effect on the percentage of cells exhibiting plasma membrane vesiculation, DNA de-condensation, nuclear rounding, DNA release into the cytoplasm or loss of cytosolic contents. However, while taxol had no effect on extracellular DNA expulsion, stabilizing the actin cytoskeleton with jasplakinolide significantly reduced the fraction of cells that expelled extracellular DNA to complete NETosis (Figure 3H, I). Examination of actin dynamics in the subset of jasplakinolide-treated cells that ruptured their plasma membrane and released extracellular DNA showed that DNA escaped through a local disruption in the cortical actin network (Movie 11). These results suggest that actin and microtubule cytoskeleton disassembly during NETosis is an active process, and that actin disassembly is important for cell membrane rupture and efficient extracellular DNA release.

### Plasma membrane permeability is progressively increased prior to its rupture and NET release

We next sought to determine how DNA escapes from the plasma membrane into the extracellular environment. We analyzed the dynamics of the plasma membrane marker CAAX-mApple in ionomycin-stimulated dHL60 cells that were either stained with SiR-DNA or co-expressing histone H2B-mEmerald to label chromatin. This showed that following microvesicle shedding, the plasma membrane appeared to remain intact through DNA de-condensation, nuclear rupture and DNA release into the cytoplasm, until a large hole was formed concomitant with extracellular DNA expulsion (figure 4A, Movie 13). This catastrophic rupture of the plasma membrane occurred much later than the previously noted decrease in contrast in the cell periphery visualized by DIC microscopy (Movie 3), suggesting that leakage of cytosol may occur by permeabilization of the plasma membrane prior to extracellular DNA release.

**Figure 4:**
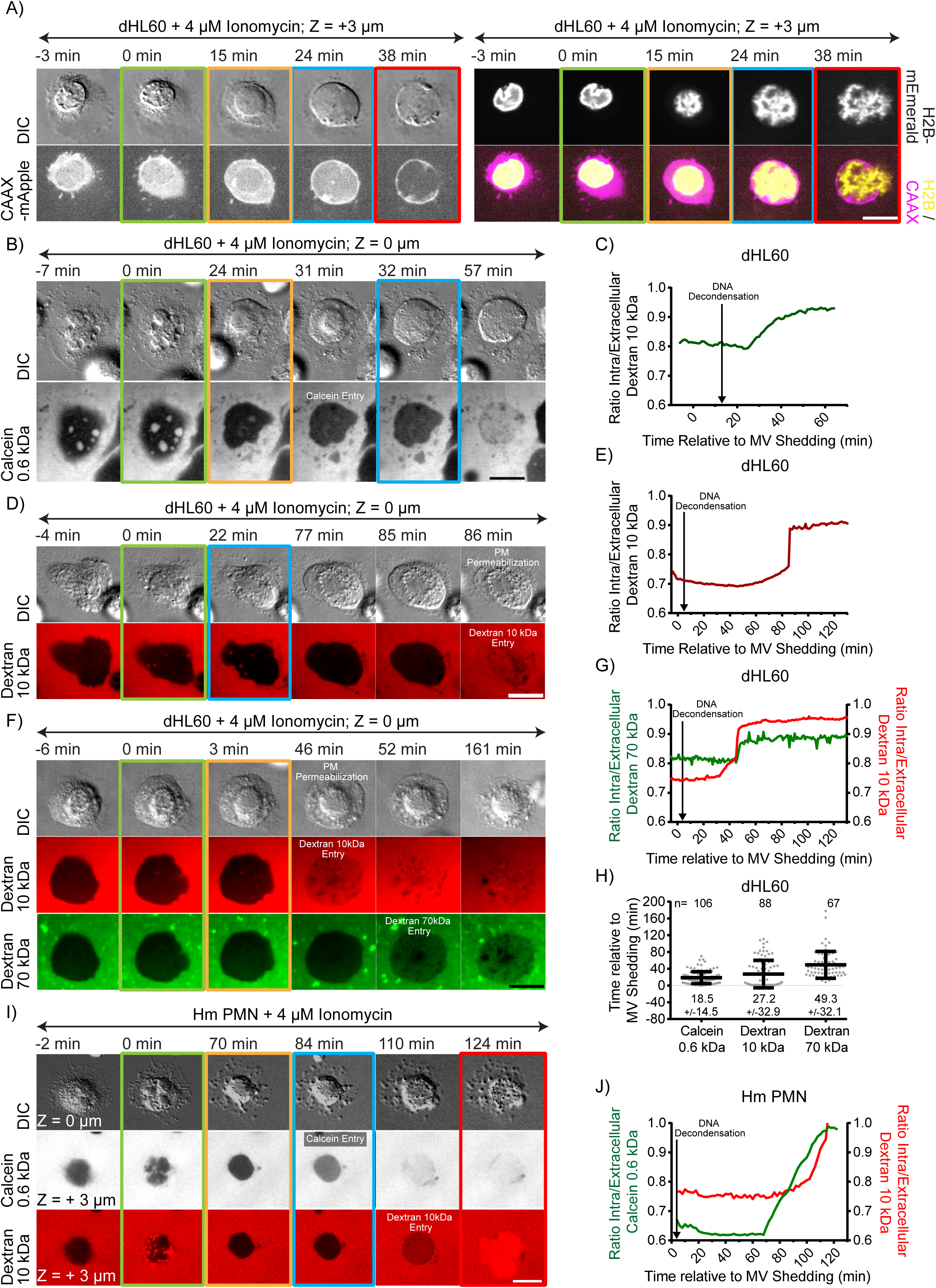
Plasma membrane becomes permeable to progressively larger molecules prior to plasma membrane rupture and NET release. Differentiated HL60 neutrophil-like cells (dHL60; A-H) or human blood polymorphonuclear neutrophils (Hm PMN; I, J) were plated on coverslips, stimulated with 4 μM ionomycin imaged live by differential interference contrast (DIC) and spinning-disk confocal microscopy at the coverslip-cell interface (Z = 0 μm) and 3 μm above in the cell center (Z = + 3 μm) at 1-2 min intervals for 4 hours, with time in min relative to plasma membrane microvesicle shedding noted above images. Double-arrow-headed lines above images indicate cells that are in presence of ionomycin, boxes around images indicate different cellular events (green: MV Shedding; yellow: DNA De-condensation; blue: DNA release to the cytosol, red: extracellular DNA release). A: Time series of DIC (upper row-Left) and fluorescence images of cell, plasma membrane (lower row-left and right, magenta), and histone (upper row-right and bottom row-right, yellow) dynamics in a dHL60 cell co-expressing the farnesylation signal sequence CAAX fused to mApple (CAAX-mApple) as a plasma membrane marker and the histones marker H2B-mEmerald. B, D, F, I: Time series of DIC (upper row) and fluorescence (lower rows) images of cell and soluble fluorescent markers (calcein (622 Da, B, I, greyscale), Alexa fluor 647 (D, F, red) or Alexa fluor 594 (I, red) 10 kDa dextran or Oregon Green 488 70 kDa (F, green)) added to the imaging media, in living dHL60 (B, D, F) or Hm PMN (I) cells. C, E, G, J: Normalized intracellular fluorescence (to average extracellular level) of calcein (C, J, green), 10 kDa (E, G, J, red) or 70 kDa (G, green) dextran over time (in min) measured from time-lapse movies. Time points of DNA de-condensation determined from the DIC channel of the movies is noted on the graphs. H: Quantification of the onset of calcein, 10 or 70 kDa fluorescent dextran cellular entry relative to microvesicle shedding determined from time-lapse DIC and confocal color-overlay movies of dHL60 cells. n = total number of cell observed, points represent individual cells, mean (long bar) and S.D. (short bar) shown below each plot. A, B, D, F, I: bars = 10 μm.

To test whether the plasma membrane became permeable prior to rupturing to expel DNA, we added different sized membrane-impermeant fluorescent markers to the imaging media and monitored their cellular entry by time-lapse spinning disc confocal microscopy of ionomycin-stimulated human or mouse PMN or dHL60 stained with SiR-DNA. We found that calcein (0.6 kDa), 10 kDa dextran, and 70 kDa dextran entered all three cell types much earlier than extracellular DNA release (figure 4B, C, D, E, H, supplementary figure 4D, E, Movie 14, 15), and mostly prior to nuclear rupture in dHL60 cells. In addition, reduction in DIC contrast in the cell periphery occurred after calcein entry and coincided with the onset of 10 kDa dextran entry (figure 4D, and F). Quantification of timing showed that calcein, 10 kDa dextran, and 70 kDa dextran entered cells on average at ∼20, ∼30, and ∼50 min after ionomycin-induced microvesicle shedding, suggesting a progressive increase in cell permeability over time as cells proceed through NETosis, Indeed, addition of either calcein together with 10 kDa dextran or 10 kDa dextran together with 70 kDa dextran showed that calcein entered cells before 10 kDa dextran (figure 4I, J), which preceded 70 kDa dextran entry (figure 2F, G, Movie 15). Similar plasma membrane permeabilization prior to NETosis was observed in mouse PMN stimulated with LPS (supplementary figure 2E, F). Together, these data show that during NETosis, the plasma membrane becomes permeable to increasingly larger macromolecules over time, allowing their entry into and leakage out from cells prior to catastrophic plasma membrane rupture and expulsion of extracellular DNA.

### PAD4 is critical to DNA de-condensation, NE rupture and extracellular DNA expulsion in mouse PMN

To further elucidate how the specific cellular events mediating NETosis depends on molecular activity, we investigated the role of the arginine deiminase PAD4, which has been shown to be required for NETosis in mouse PMN. PMN were isolated from fresh blood of wild type (WT) and PAD4 knock-out (KO) mice, stained with SiR-DNA, SiR-Actin, SiR-Tubulin or ER-dye to visualize the DNA, actin and microtubule cytoskeletons, or endoplasmic reticulum and NE, respectively, and stimulated with ionomycin. Time-lapse DIC and spinning disk confocal microscopy showed that similar to WT mouse PMN, PAD4 KO mouse PMN initiated NETosis by disassembling their actin cytoskeleton before forming and shedding plasma membrane microvesicles (supplementary figure 4F, figure 5A and B, Movie 16). In addition, WT and PAD4 KO cells both similarly vesiculated their ER networks, disassembled their microtubules, de-lobulated and rounded their nuclei and permeabilized their plasma membranes (as assessed by decreased DIC contrast and 10 kDa dextran cellular entry) (figure 5A, B, C, D, H, I, supplementary figure 4G and H, Movies 16, 17). However, in contrast to WT mouse PMN, PAD4 KO cells were deficient in DNA de-condensation, nuclear rupture, DNA release into the cytoplasm, and extracellular NET release (figure 5B, C, D, Movie 16). In addition, ER staining showed that compared to WT, the NE generally remained intact in PAD4 KO PMNs (figure 5C, I, Movie 17). Quantification of the percentage of cells that completed these events and their timing relative to microvesicle shedding showed that compared to WT, PAD4 KO displayed very strong decreases in frequency and also significant temporal delays in DNA de-condensation, nuclear rupture and extracellular DNA release in the few cells where this did occur (figure 5C, D). Immunostaining of vimentin IFs and lamin B1/B2 in WT and PAD4 KO mouse PMN at time-points after ionomycin-stimulated NETosis showed that loss of PAD4 caused delays in the disassembly of peripheral vimentin IFs (supplementary figure 4I, J) and delays in formation of lamin meshwork discontinuities (figure 5E, F, G), and blocked DNA extrusion from the nucleus. Together, these data indicate that in mouse PMN, actin and microtubule disassembly, plasma membrane and ER vesiculation, nuclear rounding and plasma membrane permeabilization occur independent of PAD4, but PAD4 is required for efficient remodeling of the vimentin IF network, DNA de-condensation, lamin meshwork and NE rupture, and extracellular DNA release.

**Figure 5:**
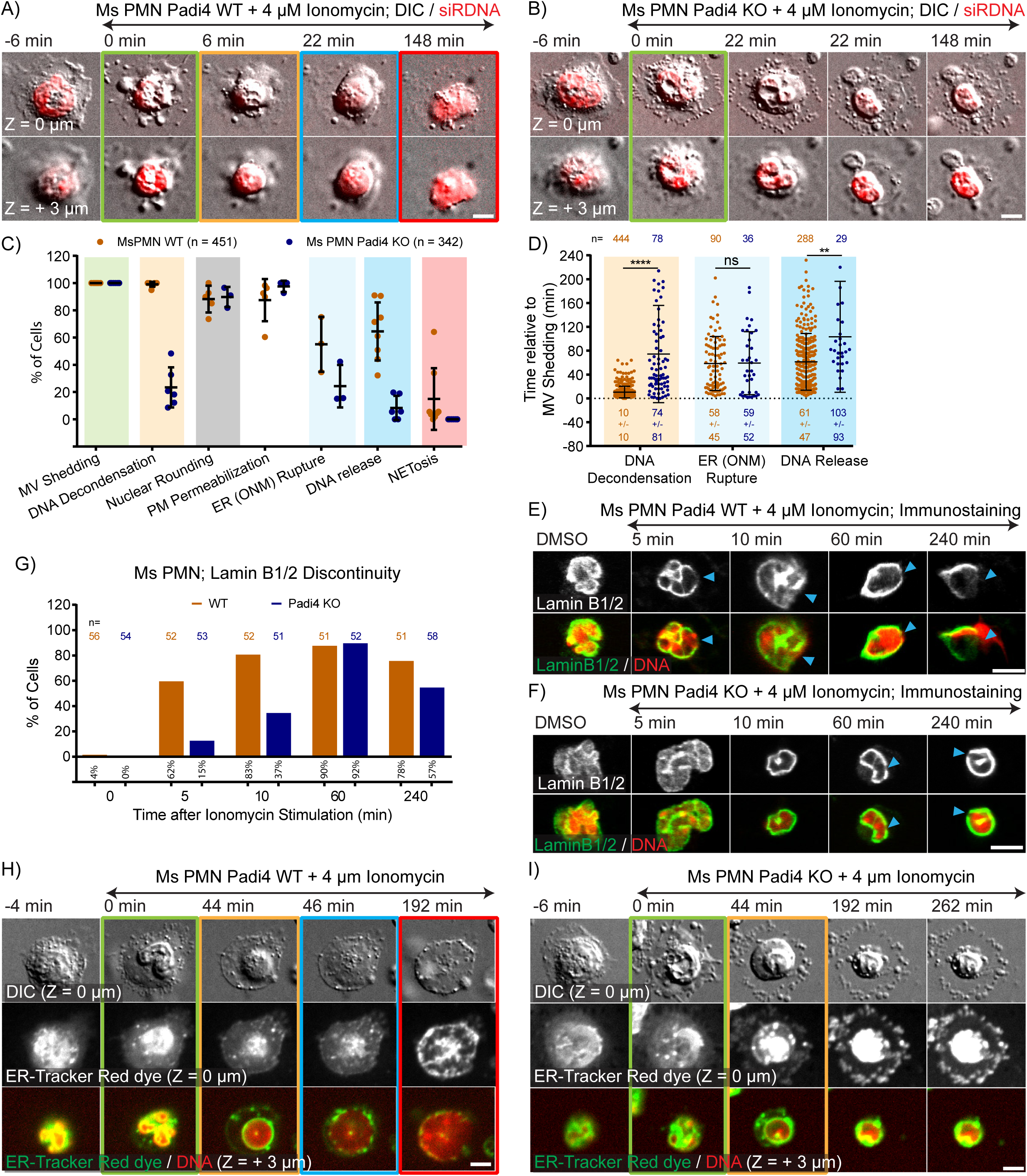
PAD4 is critical to DNA de-condensation, NE rupture and extracellular DNA expulsion in mouse PMN. Mouse blood polymorphonuclear neutrophils (Ms PMN) from wild type (Padi4 WT) or PAD4 knockout (Padi4 KO) mice were plated on coverslips, stimulated with 4 μM ionomycin, and imaged live (A, B, H, I) by differential interference contrast (DIC) and spinning-disk confocal microscopy at the coverslip-cell interface (Z = 0 μm) and 3 μm above in the cell center (Z = + 3 μm) at 2 min intervals for 4 hours, with time in min relative to plasma membrane microvesicle shedding noted above images, or fixed at different time points after stimulation (noted above images (E, F). Double-arrow-headed lines above images indicate cells that are in presence of ionomycin, boxes around images indicate different cellular events (green: MV Shedding; yellow: DNA De-condensation; blue: DNA release to the cytosol, red: extracellular DNA release). A, B: Time series of image overlays of DIC (greyscale) and SiR-DNA stained cells to fluorescently label DNA. C: Quantification of the percent of cells that exhibit microvesicle (MV) shedding, DNA de-condensation, nuclear rounding, loss of DIC contrast in the cell periphery suggesting plasma membrane permeabilization (PM Permeabilization), outer nuclear envelope rupture (ER (ONM) Rupture) quantified in cells stained with the ER-Tracker Red, release of nuclear DNA into the cytoplasm (DNA Release), and extracellular DNA release (NETosis) after ionomycin stimulation as determined from time-lapse DIC and confocal color-overlay movies. n = total number of cell observed, each point represents the percent of cells in one experiment, long bar = mean, short bars= S.D. D: Quantification of the timing relative to microvesicle shedding of DNA de-condensation, outer nuclear envelope rupture (ER (ONM) Rupture) and nuclear DNA release into the cytoplasm (DNA Release) determined from time-lapse DIC and confocal color-overlay movies. n = total number of cells observed, points represent individual cells, mean (long bar) and S.D. (short bar) shown below each plot. **** and ** for P value respectively < 0.0001, and <0.01; ns for non-significant. Statistical test: Mann-Whitney Test. E, F: Immunofluorescence of lamin B1/B2 (upper row and bottom row, green) and DAPI (at 10 min) or SiR-DNA (DMSO, 5, 60 and 240 min) staining of DNA (bottom row, red) in fixed WT (E) or PAD4 KO Ms PMN. Blue arrowheads mark rupture points in the lamina. G: Quantification from immunostaining experiments of the percent (numbers below bars) of cells exhibiting discontinuities in the lamin B1/B2 meshwork at different time points after stimulation, 0 represent cells treated with DMSO only. n = total number of observed cells. H, I: Time series of live cells stained with ER-Tracker Red (middle row and bottom row, green) to label the ER and nuclear envelope (NE) and stained with SiR-DNA (bottom row, red) showing lack of NE rupture in PAD4 KO Ms PMN. A, B, E, F, H, I: bars = 5 μm.

### PAD4 localizes predominantly to the nucleus and its enzymatic and nuclear localization activities mediate efficient DNA de-condensation, NE rupture and extracellular DNA expulsion in human dHL60

While the importance of PAD4 in mouse PMN NETosis is well documented, the role of PAD4 in human neutrophils remains controversial because of the lack of relevant model system. Patient samples with PAD4 loss-of-function mutations are not available and the currently available PAD4 inhibitors show poor efficacy in cell-based assays (46, 47) and their inhibition may be reversible (34) and thus could be incomplete. To approach the function of PAD4 in NETosis in human cells, we first characterized its localization in PMN and dHL60 cells. We performed immunostaining in dHL60 and human PMN and generated a human PAD4 cDNA fused to mEmerald (PAD4-mEmerald) and expressed it in dHL60 cells. This showed that in unstimulated (resting) dHL60 and human PMN prior to ionomycin stimulation, PAD4 primarily localized to the nucleus of both cell types as assessed by co-localization with Hoechst-stained DNA (figure 6A, B, C). In live dHL60 cells, PAD4-mEmerald co-localized with SiR-DNA or with co-expressed mCherry fused to the nuclear localization signal sequence which served as a soluble nucleoplasmic marker (NLS-mCherry) (figure 6D, Movie 18).

**Figure 6:**
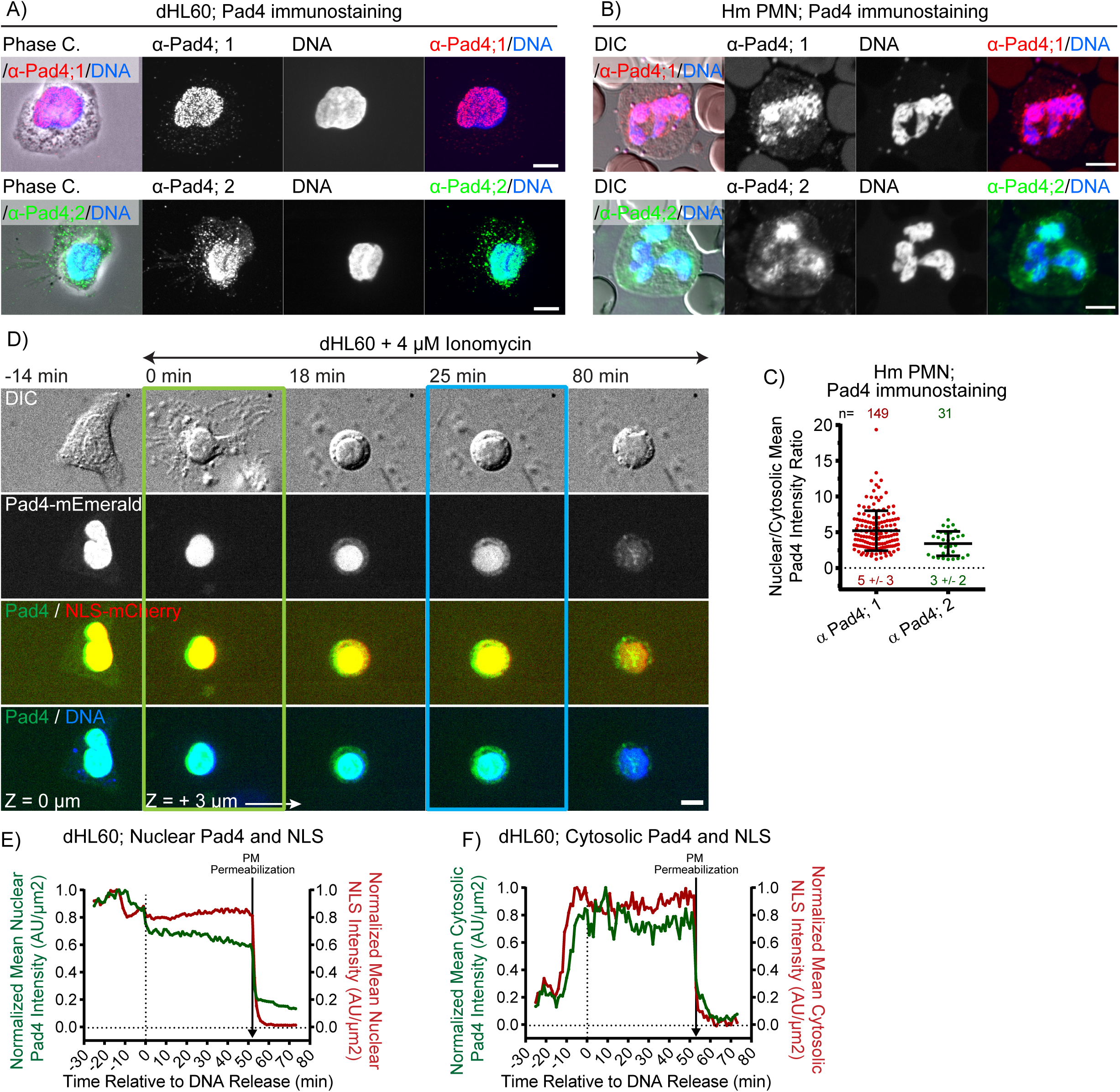
PAD4 localizes to the nucleus and enters the cytosol prior to nuclear rupture. Differentiated HL60 neutrophil-like cells (dHL60) or human blood polymorphonuclear neutrophils (Hm PMN) were plated on coverslips and fixed (A, B) or stimulated with 4 μM ionomycin and imaged by phase-contrast or differential interference contrast (DIC) and spinning-disk confocal microscopy at the coverslip-cell interface (Z = 0 μm) and 3 μm above in the cell center (Z = + 3 μm) at 2 min intervals for 4 hours, with time in min relative to plasma membrane microvesicle shedding noted above images. A, B: Phase-contrast (Phase C), DAPI staining of DNA (blue) and immunolocalization of PAD4 by two different antibodies (α-Pad4; 1 and 2, red and green) in fixed dHL60 (A) and Hm PMN (B). C: Quantification from immunostaining experiments of the mean nuclear to cytosolic ratio of PAD4 intensity using two different antibodies. n = total number of cells observed, points represent individual cells, mean (long bar) and S.D. (short bar) shown below each plot. D: Time series of DIC (top row) and fluorescence images (bottom rows) of a dHL60 cell and fluorescent PAD4 (middle row and bottom rows, green), a soluble nucleoplasmic marker (third row, red) and DNA (bottom row, blue) dynamics observed in a dHL60 cell stained with far red SiR-DNA to fluorescently label DNA and co-expressing PAD4-mEmerald and the nuclear localization signal sequence fused to mCherry (NLS-mCherry). Double-arrow-headed lines above images indicate cells that are in presence of ionomycin, boxes around images indicate different cellular events (green: MV Shedding; blue: DNA release to the cytosol). E-F: Normalized (to the maximum intensity in the compartment) mean nuclear (E) and cytosolic (F) over time (in min) of PAD4-mEmerald and NLS-mCherry measured from time-lapse movie of the cell in (D). Time point of loss of DIC contrast in the cell periphery suggesting plasma membrane permeabilization (PM Permeabilization) determined from the DIC channel of the movie is noted on the graphs. A, B, D: bars = 5 μm.

We next sought to determine the dynamics of PAD4 after stimulation of NETosis. Imaging SiR-DNA-stained dHL60 cells expressing PAD4-mEmerald after ionomycin stimulation showed that PAD4 was located primarily in the nucleus through the initial stages of NETosis including microvesicle shedding, DNA de-condensation, and nuclear rounding. However, prior to DNA release into the cytoplasm, PAD4 began accumulating more strongly in the cytoplasm, followed by a much more prominent release into the cytoplasm along with DNA that occurred concomitant with NE rupture (figure 6D, F). To determine if the accumulation of cytosolic PAD4 prior to NE rupture was a regulated export event or caused by nuclear leakage due to non-specific NE permeabilization, dHL60 cells were co-transfected with PAD4-mEmerald and NLS-mCherry, which would leak from the nucleus if NE integrity were compromised (48). Time-lapse movies showed that PAD4-mEmerald initially co-localized in the nucleus with NLS-mCherry and that the two markers simultaneously began accumulating in the cytosol after DNA de-condensation but prior to nuclear rupture (figure 6D, E, F, Movie 18). In addition, examination of DIC images showed that both PAD4-mEmerald and NLS-mCherry were rapidly released to the extracellular environment concomitant with leakage of the cytosol but before plasma membrane rupture and extracellular DNA expulsion (figure 6D, F). These results show that PAD4 in resting neutrophils is primarily localized to the nucleus. Upon stimulation of NETosis, PAD4 is first released into the cytoplasm by nuclear membrane permeabilization just after DNA de-condensation but prior to nuclear rupture. The soluble PAD4 is next released extracellularly by plasma membrane permeabilization followed by the remaining PAD4 exiting the cell along with NET expulsion.

To address the requirement for PAD4 in NETosis in human neutrophils, we took advantage of the HL60 model. As shown above, dHL60 cells exhibit a nearly identical sequence of cellular events than human PMN stimulated with ionomycin or mouse PMN stimulated with ionomycin or LPS. We generated an HL60 cell line with a modified PAD4 gene, PAD4 CR dHL60, by using CRISPR-Cas9 technology to transiently express Cas9-eGFP with a guide RNA targeting the first exon of PAD4. This resulted in near complete loss of PAD4 protein as assessed by western blot (supplementary figure 5A, B), and had no effect on HL60 differentiation into neutrophil-like cells, as assessed by their ability to produce reactive oxygen species after stimulation with PMA (20 nM, (49, 50), supplementary figure 5C, D). Stimulation of SiR-DNA-stained WT and PAD4 CR dHL60 cells with ionomycin showed that PAD4 CR dHL60 cells exhibited microvesicle shedding, nuclear rounding and plasma membrane permeabilization at a frequency and timing indistinguishable from WT (figure 7A, B, G-left). However, unlike mouse PAD4 KO cells that largely failed to de-condense their DNA, there was a less drastic reduction in the percent of cells exhibiting DNA de-condensation in PAD4 CR dHL60 cells, although the timing of this event relative to microvesicle shedding was significantly delayed compared to that in WT dHL60s (figure 7A, B, G). In addition, PAD4 CR dHL60 cells largely failed to exhibit nuclear rupture or extracellular DNA expulsion (figure 7A, B, G, Movie 19). Analysis of PAD4 CR dHL60 cells expressing Lap2β-mEmerald or laminB1-mEmerald showed that for the small fraction of cells that completed DNA release to the cytosol, both NE and lamina rupture was delayed compared to WT cells (figure 7H, I, J, K, L). Importantly, re-expressing PAD4-mEmerald in PAD4 CR dHL60 cells rescued the defects in occurrence and timing of DNA de-condensation, nuclear rupture and extracellular DNA release observed in PAD4 CR dHL60, while expressing mEmerald alone did not (figure 7C, D, G, Movie 20). Together, these data show that PAD4 is critical for efficient DNA de-condensation, nuclear lamina and NE rupture, and DNA release into the extracellular environment in both mouse and human neutrophils.

**Figure 7:**
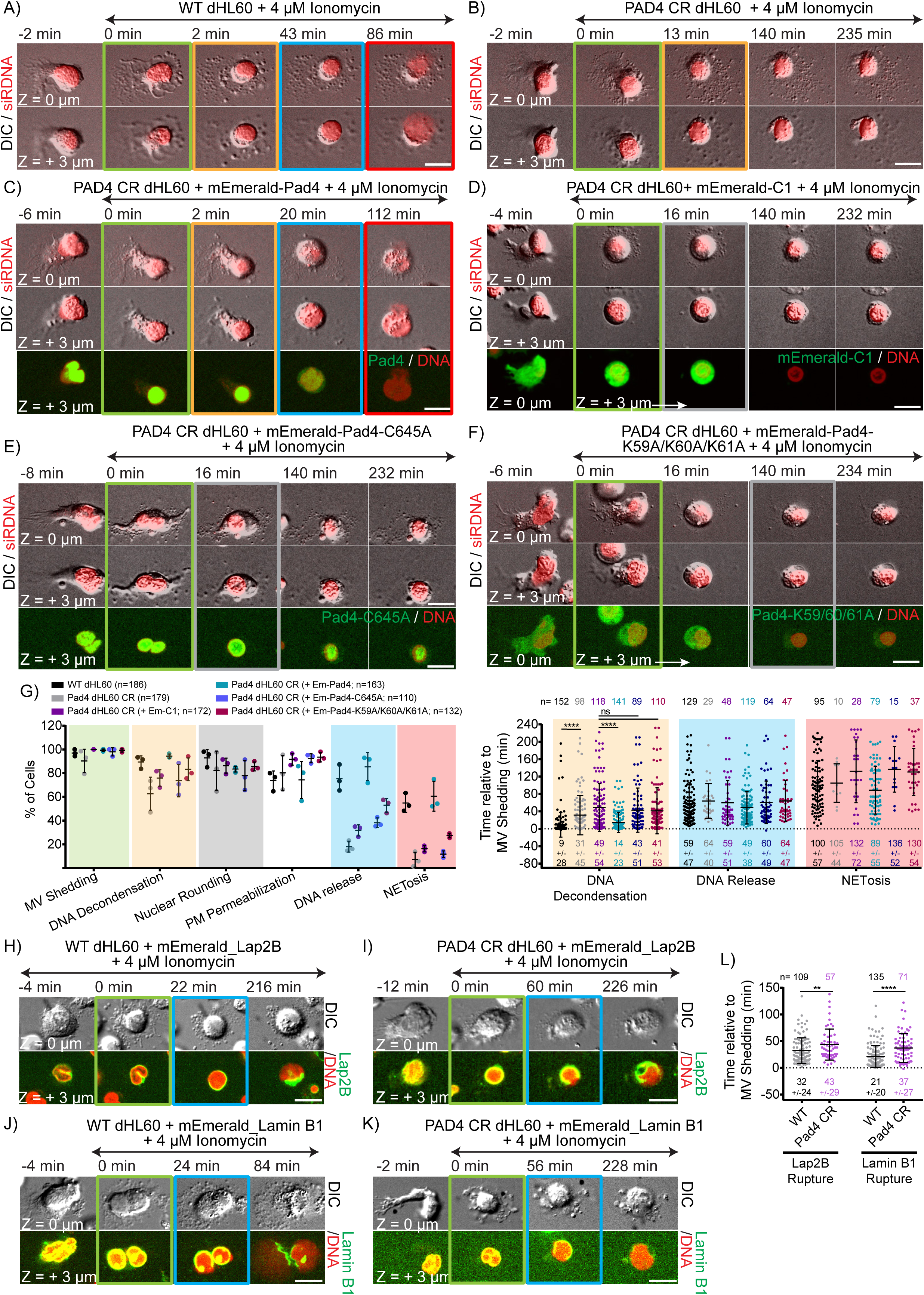
PAD4 enzymatic and nuclear localization activities mediate efficient DNA de-condensation, NE rupture and extracellular DNA expulsion in human dHL60. Differentiated HL60 neutrophil-like cells (dHL60 WT) or dHL60 with the PAD4 gene modified by CRSPR-Cas9 gene editing (PAD4 CR dHL60), were stained with far red SiR-DNA to fluorescently label DNA, plated on coverslips, stimulated with 4 μM ionomycin, and imaged live by differential interference contrast (DIC) and spinning-disk confocal microscopy at the coverslip-cell interface (Z = 0 μm) and 3 μm above in the cell center (Z = + 3 μm) at 2 min intervals for 4 hours, with time in min relative to plasma membrane microvesicle shedding noted on images. Double-arrow-headed lines above images indicate cells that are in presence of ionomycin, boxes around images indicate different cellular events (green: MV Shedding; yellow: DNA de-condensation; grey: nuclear rounding; blue: DNA release to the cytosol, red: extracellular DNA release). A-F: Time series of image overlays of DIC (greyscale) and fluorescence (red) of cell and DNA dynamics. C-F: PAD4 CR dHL60 cells were expressing mEmerald-PAD4 (C; bottom row, green), mEmerald-Only (mEmerald-C1; D; bottom row, green), enzymatically dead PAD4 mutant (mEmerald-PAD4-C645A; E; bottom row, green) or the nuclear localization mutated PAD4 (mEmerald-PAD4-K59A/K60A/K61A; F; bottom row, green). G: Quantification of the percent of cells (Left) that exhibit microvesicle (MV) shedding, DNA de-condensation, nuclear rounding, loss of DIC contrast in the cell periphery suggesting plasma membrane permeabilization (PM Permeabilization), release of nuclear DNA into the cytoplasm (DNA Release), and extracellular DNA release (NETosis) or timing relative to microvesicle shedding (right) of DNA de-condensation, DNA release and NETosis after ionomycin stimulation as determined from time-lapse DIC and confocal color-overlay movies. n = total number of cell observed, each point represents the percent of cells in one experiment (left), or individual cells (right). long bar = mean, short bars = S.D. shown below each plot (right). H-K: Time-series of DIC (upper rows) and confocal (lower rows) images of live WT or PAD4 CR dHL60 cells expressing the inner nuclear envelope protein, Lap2β-mEmerald (H, I; bottom rows, green) or lamin B1-mEmerald (J, K; bottom rows, green) and stained with SiR-DNA (bottom rows red). L: Quantification of the timing relative to microvesicle shedding of the onset of inner nuclear envelope rupture (Lap2β Rupture) or lamina rupture (LaminB1 Rupture). n = total number of cells observed, points represent individual cells, mean (long bar) and S.D. (short bar) shown below each plot. G, L: **** and ** for Pvalue respectively < 0.0001, and <0.01; ns for non-significant. Statistical test (G-left and L): Mann-Whitney Test. A-F, H-K: bars = 10 μm.

We finally sought to determine what specific activities of the PAD4 protein were required for its cellular roles in NETosis. PAD4 catalyzes arginine citrullination and is thought to mediate NETosis via this action on histones in the nucleus. To test if the citrullinating or nuclear localization functions of PAD4 were required for DNA de-condensation, nuclear rupture and extracellular DNA expulsion during NETosis, we rescued PAD4 CR HL60 cells with mEmerald-tagged PAD4 variants bearing point mutations coding amino acid substitutions that specifically inactivate these functions. PAD4 is rendered enzymatically dead by a C645A substitution (51), while its nuclear localization is disrupted by the triple K59A/K60A/K61A substitution (29, 51). Expression in unstimulated PAD4 CR dHL60 cells showed that PAD4-C645A-mEmerald localized predominantly to the nucleus (figure 7E), while PAD4-K59A/K60A/K61A-mEmerald was substantially, but not completely excluded from the nucleus prior to and after ionomycin stimulation (figure 7F), indicating that PAD4 may have an NLS- and enzymatic-independent nuclear localization mechanism in dHL60. In ionomycin-stimulated PAD4 CR dHL60 cells, expression of neither PAD4-C645A-mEmerald nor PAD4-K59A/K60A/K61A-mEmerald rescued the defects in DNA de-condensation, nuclear rupture and extracellular DNA release (figure 7C, E, F, G, Movie 20) induced by loss of PAD4. This indicates that efficient DNA de-condensation, nuclear rupture, and DNA expulsion requires PAD4 enzymatic activity in the nucleus in human neutrophils.

## DISCUSSION

Here we performed the first systematic characterization of the timing of dynamic cellular events leading to NETosis in human and mouse primary blood neutrophils, as well as in neutrophil-like dHL60 cells by using fluorescent markers of organelles and high-resolution time-lapse multi-mode microscopy. We show that NETosis proceeds by a step-wise sequence of cellular events that is conserved between mouse and human primary neutrophils, as well as in dHL60 cells, thus establishing this cell line as an important model system for *in vitro* studies of the cellular mechanisms of NETosis. For DNA to be released to the cell exterior during NETosis, it must escape from the nucleus, pass through the cytoplasm containing the dense cytoskeletons, and finally breach the plasma membrane. We demonstrate that NETosis begins with the rapid disassembly of the actin cytoskeleton, followed by shedding of plasma membrane microvesicles, disassembly and remodeling of the MT and vimentin cytoskeletons, ER vesiculation, chromatin de-condensation and nuclear rounding, progressive plasma membrane and NE permeabilization, nuclear lamin meshwork and then NE rupture to release chromatin into the cytoplasm, and finally plasma membrane rupture and discharge of extracellular chromatin (Fig 8). We find that certain cellular events always precede other events, suggesting a requirement for a precise sequence for progression through NETosis. For example, cytoskeletal disassembly, microvesicle shedding and ER vesiculation always occur before nuclear rupture, and nuclear lamina and NE rupture always precede DNA release to the cytoplasm. In contrast, plasma membrane permeabilization could occur before or after nuclear rupture but always preceded extracellular DNA release. In the three cell systems examined we found no evidence of extracellular DNA release without catastrophic rupture of the nucleus or plasma membrane or of cell viability after NET release. Using pharmacological inhibition, we also demonstrate a requirement for actin disassembly in plasma membrane rupture and extracellular DNA release. By examining cellular and organelle dynamics in mouse PMN and dHL60 human cells lacking PAD4, we demonstrate that cytoskeletal disassembly, plasma membrane and ER vesiculation, nuclear rounding, and plasma membrane permeabilization occur independent of PAD4, but PAD4 is required for efficient DNA de-condensation, NE rupture, and extracellular DNA release, and prompt timing of lamin meshwork and vimentin IF disassembly. Examination of PAD4 localization and dynamics show that PAD4 is primarily localized to the nucleus in resting neutrophils, and mutant add-back experiments show that the nuclear localization and enzymatic activities of PAD4 are critical to its role in DNA de-condensation, lamin meshwork and NE rupture, and extracellular DNA release.

**Figure 8:**
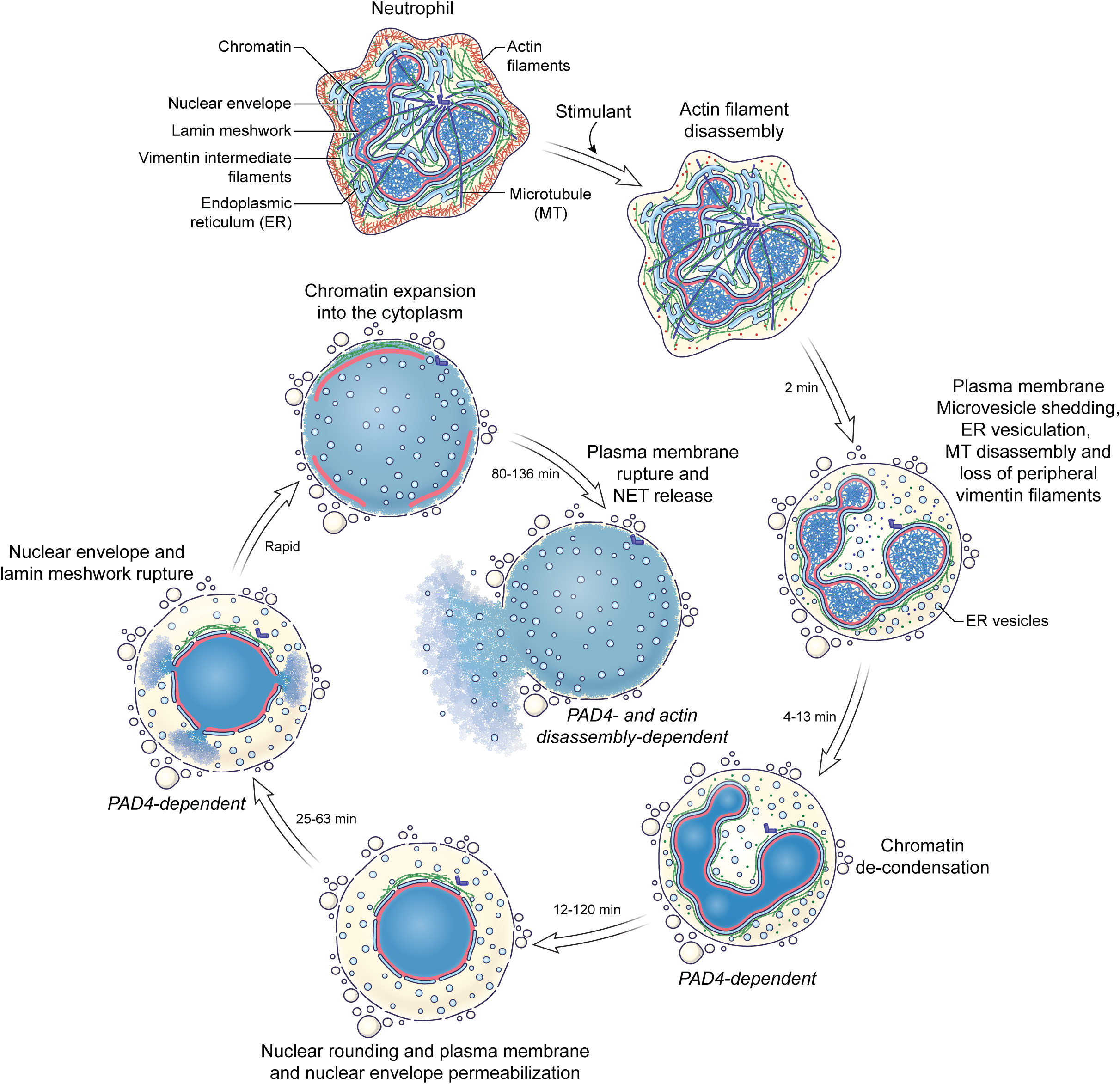
Model of the cellular events mediating NETosis. Neutrophil (first panel) refers to mouse and human PMN and differentiated HL-60 neutrophil-like cells. Times are cumulative and relative to the initiation of actin cytoskeleton disassembly, ranges indicate the range of times for all three cell types.

The conservation of cellular events between mouse and human PMNs and neutrophil-like dHL60 cells and with two different stimuli (for mouse PMNs) suggests that despite possible divergences in signaling pathways mediating the process, NETosis follows a very strict cellular mechanism. Thus, targeting these cellular pathways might be a better approach for controlling NETosis progression than targeting the various divergent signaling pathways. Indeed, plasma membrane microvesicle shedding, actin and lamin meshwork disassembly, DNA de-condensation, nuclear envelope and plasma membrane rupture are now potential therapeutic targets for NETosis inhibition.

Our study shows for the first time that the plasma membrane undergoes dramatic changes during NETosis, including shedding of microvesicles and regulated permeabilization. The shedding of microvesicles is of particular interest, as they could be important systemic messengers, providing activated neutrophils undergoing NETosis with additional ways of signaling stress (52) or contributing to disease (53). For instance, microvesicles are pro-coagulant, thus could potentially contribute to the pro-thrombotic activity of NETs (54). Of note, the microvesicles we observed at the onset of NETosis are smaller (∼<2 μm) than cytoplasts (∼4 μm, (6, 36)), and we never observed the formation of larger vesicles or cytoplasts that could be indicative of “vital NETosis”. Moreover, from a mechanical perspective, vesicle shedding is an efficient way of decreasing the plasma membrane reservoir (55, 56) thus increasing membrane tension and potentially facilitating plasma membrane rupture. In addition to microvesicle shedding, we also discovered a time-dependent, progressive increase in the size of molecules that could enter the cell prior to plasma membrane rupture and NETs release (hydrodynamic radii of calcein < 1 nm and 10 and 70 kDa dextrans are 1.9 nm and 6.8 nm respectively (57-59)). This suggests a regulated and stepwise permeabilization potentially mediated by several different membrane pores that could allow escape of specific bioactive molecules like cytokines or citrullinated peptides or proteins prior to NETs release. For example, release of PAD4 from cells prior to NETosis could enhance their prothrombotic effects, as it has recently been shown that PAD4 infusion in mice mediates ADAMTS13 citrullination, thus preventing clearance of von Willebrand factor from vessel walls and enhancing NET attachment and thrombus formation (9, 13).The mechanism of permeability is also of interest. Gasdermin D (60-63) is critical for NETosis (39, 64) and forms 20 nm pores (63), suggesting it could mediate the late stages of plasma membrane permeability during NETosis. How and why cells undergoing NETosis shed vesicles and increase their plasma membrane permeability requires further study.

Our documentation of dramatic and rapid disassembly and remodeling of the actin, microtubule and vimentin intermediate filament cytoskeletons soon after stimulation of NETosis is reminiscent of the cytoskeletal remodeling that occurs during cell division (65-67). This further supports the idea that NETosis shares similarities with mitosis (40) and suggests disassembly and/or remodeling of all three cytoskeletal systems could be mediated by mitotic kinases. We also found that the nuclear lamin meshwork undergoes disassembly during NETosis, and it has been proposed that lamin A/C phosphorylation at the same site that drives its disassembly during mitosis could occur during NETosis (Serine 22, (40, 68)). However, NETosis stimulants including ionomycin (69), PMA (70), LPS (71, 72), interleukin-8 (18) and cryptosporidium parvum (73) trigger intracellular calcium influxes. It is known that microtubules are calcium labile (74, 75), proteins capable of disassembling actin filaments including myosin II and gelsolin can be calcium-dependent (76-78), and kinases and proteases regulating vimentin disassembly or degradation can be activated by calcium (79-82). Hence, it is possible that cytoskeletal remodeling and disassembly during NETosis may be mediated by calcium. Our finding that actin disassembly was required for plasma membrane rupture and extracellular NET release indicates that molecules mediating actin disassembly could serve as therapeutic targets for NETosis inhibition.

Our study provides the first systematic analysis of the role of PAD4 in the cellular events of NETosis using genetically modified mouse PMN and differentiated HL60 neutrophil-like human cells. This approach precludes problems associated with PAD4 inhibitors including lack of specificity due to activity against other PAD family members or lack of reversibility (34, 46, 47). The role and requirement of PAD4 for NETosis has been controversial particularly regarding human neutrophil NET release. For instance, histone citrulliation by PAD4 has been proposed to be only a consequence of NETosis and not crucial for the process (24). However our results show that both the citrullination activity and nuclear localization signal of PAD4 are required for NETosis, strongly supporting the notion that PAD4 enzymatic activity in the nucleus, potentially through histone citrullination (23) mediates NETosis. Although PAD4 is critical for DNA de-condensation, lamin meshwork and NE rupture and extracellular DNA release in mouse PMNs and HL60-derived neutrophils, subtle differences in the extent of PAD4 involvement exist. For example, the role of PAD4 in DNA de-condensation appears stronger in mouse PMN than in dHL-60 cells. Although this could be attributed to the residual (∼5%) PAD4 in the gene-edited HL-60 cell line, it is also possible tthat neutrophil enzymes elastase, MPO, PR3 or cathepsin G may be more important for de-condensation in human than in mouse cells (25). Despite the differential effect of PAD4 on DNA de-condensation between mouse PMN and dHL60 cells, PAD4 depletion had similar negative consequences on release of DNA from the nucleus into both the cytoplasm and extracellularly, indicating that full de-condensation is required for DNA release. Finally, since PAD4 depletion delays the disassembly of lamins and vimentin, the two sets of intermediate filaments involved in the maintenance of nuclear mechanical integrity (83-85), it is tempting to speculate that PAD4-mediated citrullination of these proteins could drive nuclear envelope rupture during NETosis, perhaps by promoting their disassembly. The cellular targets of PAD4 that are required for NETosis remain to be determined.

## ACKNOWLEDGEMENTS

The authors wish to thank the Marine Biological Laboratory for support through the Whitman Fellows program (DDW), Nikon Instruments for loan of microscopy equipment (CMW) and support through the Nikon Fellows program (DDW) at the Marine Biological Laboratory, Chloe Bulinski, James Anderson and the late Mike Davidson for the gift of reagents, Schwanna Thacker for administrative assistance, William Shin for technical support with microscopy, Robert Fischer and Ana Pasapera for helpful suggestions, the NHLBI flow cytometry core facility, and the NHLBI Blood Bank. This work is funded by the NHLBI Division of Intramural Research (CMW and HRT), NIH RO1 GM106023 and NIH PO1 GM096971 (RDG, AEG, RQ, MK and AV), NIH T32 CA08062160038140 (MK and AV), and NIH R35 HL135765 (DDW and SLW).

## SUPPLEMENTARY MATERIALS

### MATERIALS AND METHODS

#### Cells

##### HL-60 cell differentiation and transfection

HL-60 cells were purchased form ATCC (ATCC^®^ CCL-240^™^), cultured at 37°C and 5% CO_2_ in RPMI 1640 with L-glutamine (ThermoFisher, 11875093) with 25 mM HEPES (ThermoFisher, 15630080), 1% Penicillin and Streptomycin (P/S; ThermoFisher, 10378016) and 15% heat-inactivated Fetal bovine Serum (FBS; Atlanta Biologicals, S11150) and split every 2 days at 3.10^5^ cells/mL. Cells were differentiated with culture media supplemented with 1.3% DMSO (Sigma, D2650) as previously described (1). Differentiated cells (dHL60) were used 6 or 7 days after onset of differentiation. Transient expression of cDNA was performed by electroporating 2 µg of DNA in 2×10^6^ cells using the Amaxa nucleofector kit V (Lonza, VCA-1003) and the program Y-001 on an Amaxa Nucleofector II (Lonza). Differentiated cells were imaged 6 hours after transfection. HL60 cells stably expressing F-tractin-mApple were generated by transfecting 2×10^6^ non-differentiated HL60 cells with 2 µg of DNA. Transfected cells were selected with 1 mg/mL of G418 for 2 weeks before being sorted by flow cytometry for mApple positive cells. All live cell imaging was performed with imaging media: RPMI 1640 lacking Phenol red (Fisher scientific, 11835030) with 25 mM HEPES and 1% P/S.

##### Isolation of mouse neutrophils

Neutrophils were isolated from the fresh blood of WT and PAD4 KO mice as previously described (2). Briefly, the mice were bled for 1 mL via the retro-orbital plexus into 2 mL anti-coagulant (15 mM EDTA and 1% BSA in sterile PBS) after being deeply anesthetized with isoflurane. The blood was centrifuged at 2,400 rpm for 12 min at room temperature, the supernatant was removed, and cells were resuspended in anti-coagulant before loading on top of a percoll gradient column of 52%/69%/78%, with 78% being the lowest layer in the 15 mL centrifuge tube. The column was then centrifuged at 1,500 x g for 32 min at room temperature with acceleration set at 3 and deceleration set at 0. Cells at the 69%/78% interface were collected and pelleted with addition of PBS at 500 x g for 12 min at room temperature. Cells were then subjected to a brief water lysis, and the majority of the red blood cells were removed after the osmotic challenge. The neutrophils were then resuspended in imaging media, either stained with SiR-DNA, SiR-Actin, SiR-Tubulin or ER-Tracker Red dye according to manufacturers’ directions for immediate live imaging, or plated onto coverslips and stimulated for NETosis, and thereafter fixed for immunostaining.

##### Isolation of human blood neutrophils

Human blood samples were obtained from healthy donors on NIH IRB-approved protocol 99-CC-0168. Research blood donors provided written informed consent and blood samples were de-identified prior to distribution. Neutrophils isolation was performed as described in (3). Briefly, 6 mL of blood was layered on 6 mL of Histopaque-1119 (Sigma-Aldrich, 11191) then spun down at 1100 g for 22 min at 24°C; acceleration 2, deceleration 1. After removal of the plasma and leukocyte layer, the granulocytes/erythrocytes layer was washed with HBSS (Gibco™ HBSS without Calcium, Magnesium or Phenol Red; ThermoFisher, 14175095) by centrifugation (400 g; 10 min; 23°C; acceleration 9; deceleration 9). The resulting pellet was resuspended in 2 mL of HBSS then layered on a percoll (Percoll Plus, GE Healthcare, 17-5445-02) gradient column of, from bottom to top, 85%/80%/75%/70%/65% percoll in PBS (DPBS, no calcium, no magnesium; ThermoFisher, 14190144) before being centrifuged at 1100 g for 23 min at 23°C; acceleration 2, deceleration 1. The neutrophil layer (from the middle of the 70% to the middle of the 80% percoll layer) was washed with HBSS by centrifugation (400 g; 11 min; 23°C; acceleration 9; deceleration 9). The pellet was resuspended in 1 mL HBSS before being counted. Cells were resuspended at 10^3^ cells/µL in imaging media and 3×10^5^ cells were either stained with SiR-DNA, SiR-Actin, SiR-Tubulin or ER-Tracker Red dye according to manufacturers’ directions for live cell imaging or plated on coverslips for immunostaining.

#### cDNA Expression Vectors

cDNAs encoding CaaX-mApple, H1-mEmerald, ER-5-mEmerald, Lamin B1-mApple, Lamin A-mEmerald, F-tractin-mApple, G-actin-mEmerald, Tubulin-mEmerald, Vimentin-mEmerald, H2B-mEmerald and mEmeral-C1 were the kind gift of the late Mike Davidson (Florida State University, Tallahassee, FL); eGFP-Ensconsin (3X-GFP-EMTB) was a kind gift from Chloe Bulinski (Columbia University, New York, NY); pSpCas9-2A-GFP vector (Addgene, 48138 (4)) was a kind gift from James Anderson’s Lab (NHLBI/NIH, Bethesda, MD).

To generate the PAD4-mEmerald vector, human PAD4 cDNA was purchased from Sino Biological Inc. (pCMV3-N-His-PADi4, HG11072-NH). The PADi4 mRNA was PCR amplified with primers containing the KpnI and BamHI restriction sites (Fw1: 5’-CCG GGT ACC GCC CAG GGG ACA TTG ATC CG-3’; Rev1: 5’-GGT GGA ACA TGG TGC CCT AAG GAT CCC CG-3’). The so obtained PAD4 insert and the mEmerald-C1 vector were cut with restriction enzymes (KpnI, BamHI, New England Biolabs) then ligated together with T4 DNA ligase (ThermoFisher, EL001) after dephosphorylation with Calf Intestinal Alkaline Phosphatase (CIAP; ThermoFisher, 18009019) of the linearized plasmid and gel purification (QIAquick Gel Extraction Kit; Quiagen, 28706) of both insert and vector. DH5α competent cells (ThermoFisher, 18258012) were transformed with 5 µL of the ligation product. Kanamycin (ThermoFisher, 15160054) resistant colonies were further amplified and minipreps (Quiagen; 27106) were performed. The obtained mEmerald-PAD4 plasmids were cut with the restriction enzymes KpnI and BamHI to verify the presence of a 2 kbp and 4.7 kbp products testing the overall integrity of the plasmid. The insert was further verified by full sequencing.

The enzymatically dead (mEmerald-PAD4-C645A) and the NLS dead (mEmerald-PAD4-K59A/K60A/K61A) mutants were generated based on (5) and (6) respectively by *in vitro* site-directed mutagenesis using the QuikChange II XL Site-Directed Mutagenesis Kit (Agilent, 200521) as directed by the manufacturer. Briefly, appropriate mutations were introduced to the mEmerald-PAD4 vector by PCR amplification using following primers: K59-60-61A-Fw: 5’-CCA TGT GGA GGA ACC TGT GGA TGC CGC CGC GGC TGG AGG GCC GTG GGC AAT A-3’; K59-60-61-Rev: 5’-TAT TGC CCA CGG CCC TCC AGC CGC GGC GGC ATC CAC AGG TTC CTC CAC ATG G-3’; C645A-Fw: 5’-CAC GTT GGT GCC GGC GTG CAC CTC CCC A-3’; C645A-Rev: 5’-TGG GGA GGT GCA CGC CGG CAC CAA CGT G-3’. The PCR amplification products were incubated with DpnI restriction enzyme to digest the parental methylated and hemimethylated DNA (mEmerald-PAD4) before being used to transform XL10-Gold Ultracompetent Cells. Kanamycin resistant colonies were further amplified and minipreps were performed. The presence of the desired mutations was confirmed by sequencing.

#### Inhibitors

Jasplakinolide (J7473) and Taxol (Paclitaxel; P3456) were purchased from Thermo Fisher Scientific.

#### Generation of gene-edited HL60 PAD4 Crispr cell line

The generation of the PAD4 Crispr line was performed as described (4). The PAD4 guide RNA was designed using the sequence of the first exon of the human PAD4 gene and the Crispr.mit webtool from the Zhang Lab (http://crispr.mit.edu). The following guide RNA sequence was selected: Guide1: 3’->5’ TC ACA CGG ATC AAT GTC CCC TGG. The selected guide RNA had a low score (<0.3) of possible exonic off-targets with more than 4 mismatches per sequence. The following oligos containing the guide RNA without the PAM sequence but with the 5’-CACC and the 5’-AAAC overhangs were used: Hm-Padi4-Exn1-1-Fw: 5’-CAC CGG GGA CAT TGA TCC GTG TGA-3’; Hm-Padi4-Exn1-1-Rv: 5’-AAA CTC ACA CGG ATC AAT GTC CCC-3’.

Oligos were phosphorylated with the T4 Polynucleotide Kinase (NEB, M0201S), annealed then ligated into the pSpCas9-2A-GFP vector using the T4 DNA ligase. The bacterial genomic DNA was removed using the Plasmid Safe exonuclease (Fischer Scientific, NC9046399) then the resulting sgRNA-pSpCas9-2A-GFP vector was transformed into One Shot^®^ Stbl3™ Chemically Competent *E. coli* (Thermo Fischer, C737303). Ampicillin resistant colonies were further amplified, plasmid DNA was purified using the QIAprep Spin Miniprep Kit and the insertion and integrity of the guide RNA sequence was verified by sequencing using the LKO1 primer (5’-GAC TAT CAT ATG CTT ACC GT-3’). Validated sgRNA-pSpCas9-2A-GFP plasmid were amplified and purified using the PureYield™ Plasmid Maxiprep System (Promega, A2393) to generate pSpCas9-2A-GFP_Padi4-sgRNA.

5×10^6^ non-differentiated HL60 cells at passage 5 (passage 0 being when cells were received from ATTC) were electroporated with 5 µg of the pSpCas9-2A-GFP_Padi4-sgRNA plasmid using the Amaxa nucleofector kit V and the program Y-001 in an Amaxa Nucleofector. 24 hours after transfection, GFP positive cells were sorted by flow cytometry into 348 well plates (1 cell/well). Clonal cells were expanded and differentiated with DMSO and PAD4 protein level verified by western blot. All the obtained 20 clones had some remaining PAD4 protein detectable by western blot. The clone with the lowest level of PAD4 protein was retained and used as the HL60 PAD4 CR line.

#### Immunofluorescence

##### Lamin and vimentin immunostaining of mouse PMN

Neutrophils freshly isolated from the blood of WT and PAD4 KO mice were resuspended in serum-free RPMI 1640 medium supplemented with 25 mM HEPES and 1% P/S and plated on cleaned coverslips (#1.5, 22 mm x 22 mm) on top of parafilm in 35 mm wells at 2.1 × 10^4^ cells in 100 μL per coverslip. Cells were incubated at 37°C for 5 min to allow adherence before the addition of ionomycin (4 μM; Tocris, 1704) or vehicle (DMSO). After 4 hours, cells were fixed and permeabilized with anhydrous −20 °C methanol (Alfa Aesar 41838AK) for 10 minutes. Fixed cells were rinsed twice with 0.05% TWEEN 20 (Sigma, P1379) for 3 minutes each, followed by a third wash with phosphate buffered saline (PBS, Gibco, ThermoFisher, 20012027) for another 3 minutes, using 2 mL for each wash. Cells were then incubated with primary antibodies (rabbit anti-LB1/2 (1:1000; Abcam ab16048) or chicken anti-vimentin (1:250; Biolegend Poly29191)) for 1 hour at room temperature and rinsed again as described above. Next, samples were incubated with secondary antibodies (Alexa 488 goat anti-rabbit, 1:400, ThermoFisher, A-11008; or Alexa 568 goat anti-chicken, 1:400, ThermoFisher, A-11041) for an hour at room temperature and stained with either Hoescht 33258 (1:10000, Molecular Probes, H3569) or SiR-DNA (1 µM, Cytoskeleton, CY-SC007) and rinsed again. Stained cells were then mounted in Prolong Glass (ThermoFisher, P36982) and allowed to cure for 24 hours. Gentle vacuum suction was used between each step above to remove the solution from the prior step.

##### HL60 and Human PMNs lamin and vimentin immunostaining

3×10^5^ dHL60 cells or Human PMNs were resuspended in 300 µL of culture media without FBS (RPMI 1640, 25 mM HEPES and 1% P/S) and plated on 22 × 22 mm #1.5 cleaned glass coverslips. Cells were left to adhere on coverslips for 10 minutes at 37°C and 5% CO_2_. For Human PMNs, 700 µL of culture media without FBS supplemented with 1 µL of DMSO or Ionomycin (4mM; for a final concentration of 4 μM) was added to coverslips then incubated for 5, 10, 60 or 240 minutes at 37°C and 5% CO_2_. Indirect immunofluorescence was performed as already described (7). Briefly, cells were fixed for 20 minutes at 37°C with 4% paraformaldehyde (PFA; Electron Microscopy Science,15710) in cytoskeleton buffer (CB; 10 mM MES, 3 mM MgCl2, 138 mM KCl, and 2 mM EGTA) then permeabilized with 0.5% of Triton X-100 in CB at 37°C for 5 minutes. Free aldehydes were quenched with 10 mM glycine in CB at room temperature (RT). Cells were washed once for 5 minutes followed by 2 × 10 minutes washes with Tris Buffered Saline (TBS; 20 mM Tris, pH 7.6, 137 mM NaCl2) before being blocked for 1 hour at RT with blocking solution (2% BSA IgG free and protease free (Sigma-Aldrich, A3059); 0.1 % Tween 20 (Sigma-Aldrich, P1379) in TBS). Cells were stained for 2 hours at RT with primary antibodies (1:400) diluted in blocking solution then washed before being incubated with fluorophore-conjugated secondary antibodies (1:200), Alexa-Fluor-647 phalloidin (1:100; ThermoFisher, A22287) and Dapi (1ug/mL; Sigma, 268298) diluted in blocking solution, for 1 hour at RT. Cells were washed with blocking solution then with TBS (2 × 10 minutes, each). The coverslip was mounted on a glass slide in mounting media (Dako; Pathology Products, S3023) then sealed with nail polish. The following primary antibodies were used: rabbit anti-lamin A/C (abcam, ab8984), rabbit anti-lamin B1 (abcam, ab16048), mouse anti-lamin B2 (abcam, ab8983), mouse anti-PAD4 (abcam, ab128086), rabbit anti-PAD4 (GeneTex, 113945), mouse anti-vimentin (Clone 091D3, Biolegend, 677801). The secondary antibodies Alexa Fluor 488 Donkey anti-rabbit (711-545-152), Alexa Fluor 594 Donkey anti-rabbit (711-585-152), Alexa Fluor 488 Donkey anti-mouse (715-545-150) and Alexa Fluor 594 Donkey anti-mouse (715-585-150) were from Jackson Immunoresearch Laboratories.

##### PAD4 immunostaining of human PMNs

Fresh blood was collected from 3 human subjects into EDTA-coated vacutainers or capillaries, and 3-5 μL was immediately smeared onto microscope slides. The slides were allowed to air dry for 2-3 min, and then fixed in 100% methanol for 10 sec. After air dry for another 2-3 min, cells were fixed in 2% paraformaldehyde for 2 hours at room temperature. The slides were washed 3 times with PBS and permeabilized in 0.1% sodium citrate and 0.1% Triton X-100 in PBS for 10 min at 4°C. After a rinse in PBS, the slides were blocked in 3% BSA in PBS for 2 hr at room temperature in a humidified chamber. Primary antibodies diluted in 0.3% BSA with 0.05% Tween-20 in PBS (PBS-T-BSA) were added to the slides: (1) anti-PAD4 (1:500; Abcam, ab128086) plus anti-CD11b (1:250; Abcam, ab52478) or (2) anti-PAD4 (1:100; GeneTex, 113945) plus anti-CD66b (1:250; Biolegend, 305103), and incubated overnight at 4C. After two washes in PBS, appropriate Alexa Fluor-conjugated secondary antibodies were added to the slides at 1:1,500 diluted in PBS-T-BSA. The slides were then washed, counterstained for DNA using Hoechst 33342 (1:10,000) for 10 min at room temperature, and coverslips were mounted with FLUORO-GEL mounting medium (Electron Microscopy Sciences).

#### dHL-60 NETosis endpoint assay

Differentiated HL60 cells were pelleted, resuspended in serum-free RPMI 1640 medium buffered with 10 mM HEPES, and plated at 15,000 cells per well in 96-well plates. After 20 min of incubation to allow for adherence, cells were stimulated with ionomycin (4 μM) for 2.5 h, then fixed in 2% paraformaldehyde containing Hoechst 33342 (1:10,000; Invitrogen, H3570) at 4°C overnight. Fixed cells were imaged on a Zeiss Axiovert 200M wide-field fluorescence microscope. Percentage of NETs was quantified from 6 non-overlapping fields per well and the average was taken from triplicates in every experiment.

#### Western blot

Western blots were performed as previously described (8). In brief, cells were lysed in Laemmli sample buffer, lysates were separated by SDS-PAGE and proteins were electro transferred overnight at 4°C to an Immobilon-P PVDF membrane. Membranes were blocked for 1 h at room temperature (RT) with 5% nonfat dry milk (wt/vol) in TBS-T buffer (TBS + 0.1% Tween-20 [vol/vol]) then incubated for 2 h at room temperature with indicated primary antibodies. Subsequently, membranes were washed 3 × 5 min in TBS-T, incubated with appropriate HRP-conjugated secondary antibodies (1:10,000) for 1h at room temperature then washed 3 × 5 min in TBS-T. An ECL detection system (Millipore) was used to visualize protein bands.

The following antibodies were used: mouse anti-PAD4 (1:1000; abcam, ab128086), rabbit anti-lamin A/C (1:1000; abcam, ab8984), rabbit anti-lamin B1 (1:1000; abcam, ab16048), mouse anti-lamin B2 (1:1000; abcam, ab8983), rabbit anti-Gapdh (1:2000; Clone 14C10, Cell Signaling, 2118S). The secondary antibodies (HRP-conjugated goat anti-mouse (115-035-003) or goat anti-rabbit (111-035-003)) were from Jackson ImmunoResearch Laboratories.

The relative level of PAD4 protein in CRSPR cell lines was quantified using ImageJ 1.52n (NIH) by first performing local background subtraction around the bands of interest, then calculating the ratio between the PAD4 intensity and the corresponding Gapdh intensity for both WT and PAD4 CR dHL60 cells to obtain the PAD4 protein amount relative to the total protein content. For each repeat, the so normalized PAD4 intensity was further normalized to the WT dHL60 level to obtain the percentage of PAD4 protein relative to WT.

#### Nitroblue tetrazolium (NBT) assay for reactive oxygen species production

The NBT assay was performed as described (9, 10). 2×10^6^ cells were washed two times in PBS at room temperature then resuspended in 1.5 mL of imaging media alone(-PMA; -NBT) orsupplemented with 20 nM of phorbol-12-myristate-13-acetate (PMA) (+PMA; -NBT), 3 mg of NBT (ThermoFisher, N6495) (-PMA; +NBT), or 20 nM of PMA combined with 3 mg of NBT (+PMA; +NBT). Cells were then incubated for 25 minutes at 37°C and 5% CO_2_ before being transferred on ice, washed three times in ice cold PBS then fixed with 100 µL of ice cold 2% PFA in PBS. Fixed cells were left overnight at 4°C before being analyzed by flow cytometry.

Flow cytometry was performed on an LSR II equipped with a BD FACSDiva™ software (BD Biosciences). Cells’ side scatter area (SSC-A) as function of cells’ forward scatter area (FSC-A) that measure cell granularity and size respectively plots were generated. Control cells (-PMA; - NBT) were first loaded to define the NBT<0 gate on the FSC/SSC plot and the new population with higher SSC-A detected by loading the (+PMA; +NBT) sample was used to define the NBT>0 gate. Samples were then run at medium flow rate and detection was stopped at 100.000 cells. The percentage of cells in the NBT<0 and NBT>0 was measured with the FACSDiva™ software.

#### Reverse Transcriptase Polymerase Chain Reaction (RT-PCR) for PADi4 mRNA detection

RNA from differentiated WT and PAD4 CR HL60 cells was extracted using the RNeasy Mini Kit (Quiagen, 75104) as directed by the vendor. RT-PCR was performed using the AccessQuick™ RT-PCR System (Promega, A1701) as directed by the vendor with Gapdh as loading control. The following primers were used: PAD4 P1 Fw: 5’-ACT TCT TCA CAA ACC ATA CAC TGG-3’; PAD4 P1 Rev: 5’-CCT CGA GTT ACA TAG CCA AAA TCT-3’; PAD4 P2 Fw: 5’-GTG TTC CAA GAC AGC GTG GT −3’; PAD4 P2 Rev: 5’-GTT TGA TGG GAA ACT CCT TCA G-3’; GAPDH Fw: 5’-ACC CAG AAG ACT GTG GAT GG-3’; GAPDH Rev: 5’-CCC CTC TTC AAG GGG TCT AC-3’. The PCR product was separated by electrophoresis using a 1% agarose gel in Tris Acetate EDTA buffer stained with Midori green advance (1:20,000; Nippon Genetics, MG04). mRNA bands were visualized using the Ultraviolet mode of MyECL imager (ThermoFisher).

#### Microscopy

##### Spinning disc confocal and DIC microscopy

Imaging was performed on a Nikon Eclipse Ti or Ti2 microscope equipped with Perfect Focus™ equipped with a Yokogawa CSU-X1 or CSU-W1 spinning disc scanhead, a Hamamatsu Orca-flash 4.0 v2 or v3 camera and a Plan Apo 60x oil 1.4 NA DIC, an Apo Tirf 60x oil 1.49 NA DIC or an Apo Tirf 100x oil 1.49 NA DIC Nikon objective lens and the appropriate DIC prisms in place. Illumination was provided on the CSU-W1 by Nikon LUNV 6 line laser unit (20 mW 405nm; 20 mW 445nm; 70 mW 488nm; 40 mW 515nm; 70 mW 561nm and 40 mW 640nm; power measurement is at the fiber output) with 2 single mode optical fiber outputs (1 APC and 1 UPC on a fast actuator). Illumination on the CSU-X1 was provided by the Agilent MLC400B Monolithic Laser Combiner with four lasers (405nm: 20mW, 488nm: 50 or 80mW, 561nm: 50 or 80mW, 647nm: 125mW)). DIC illumination was provided by a 100W halogen bulb or LED using an 0.52 NA condenser lens. Microscopes were equipped with the Nikon motorized stage with xy linear encoders and a Mad City (Madison, WI) Nano-Z100 piezo insert with 200um travel. Laser confocal or DIC illumination were selected with electronic shutters and an automated filter turret containing a multibandpass dichromatic mirror together with an electronic emission filterwheel. Microscope functions were controlled by NIS-Elements software (Nikon).

For live cell imaging, cells were incubated with SiR-DNA (30 min at 0.5 or 1 µM; Cytoskeleton, CY-SC007), SiR-Actin (1 hour at 1 µM; Cytoskeleton, CY-SC001), SiR-Tubulin (1 hour at 1 µM; Cytoskeleton, CY-SC006) or ER-Tracker Red dye (30 min at 1 µM; ThermoFisher, E34250) at 37°C and 5% CO_2_ before being washed then resuspended in 300 µL of imaging media. For assaying plasma membrane permeability cells were imaged in media containing 20 µM calcein (ThermoFisher, C481), 1 mg/mL of Alexa fluor 647 (ThermoFisher, D22914) or Alexa fluor 594 (ThermoFisher, D22913) 10 kDa dextran or Oregon Green 488 70 kDa dextran (ThermoFisher, D7172). A non-coated, gamma-irradiated glass bottom 35 mm dish (WPI, FD35-100), 24-well plate (CellVis, P24-1.5H-N) or 4 Well chambered cover Glass (CellVis, C4-1.5H-N) was placed on a pre-warmed (37°C) microscope stage. Sample temperature and humidity were controlled with a Tokai Hit stage-top incubator (Tokai Hit) or a Pathology devices live cell stage-top incubator (Pathology Devices). Cells were plated on temperature-equilibrated glass bottom dishes and allowed to adhere for 5 minutes. Multiple (3 to 10) random fields containing multiple adherent cells were selected for imaging. Confocal and DIC images at the coverslip-cell interface and 3 µm above in the cell center were acquired for each position. The image acquisition was paused 10 minutes after beginning, ionomycin (final concentration of 4 µM) or lipopolysaccharide (LPS (Klebsiella pneumoniae) final concentration of 25 µg/mL; Sigma-Aldrich, H4268) was added and imaging was resumed. Images sets of DIC, 488 nm (Calcein,, Oregon Green dextran, EGFP and mEmerald excitation), 561 nm (AlexaFluor 594 dextran, mApple and ER-Tracker Red dye excitation) and 647 nm (Alexafluor 647 dextran, SiR-DNA, SiR-Actin and SiR-Tubulin excitation) captured in rapid succession were acquired every 1-2 minutes for 4 hours.

dHL60 and Human PMNs that were immunostained for lamins and vimentin were imaged by spinning disc confocal microscopy with a Plan Apo 100x oil 1.4 NA Ph3, DM Nikon objective and a 0.85 NA Dry condenser. For each coverslip, multiples (3-6) random fields were selected and pairs of phase contrast and fluorescence confocal stacks were captured.

##### Laser scanning confocal microscopy

Mouse PMNs that were immunostained for vimentin and laminB1/2 were imaged using a Zeiss LSM 510 laser scanning confocal microscope (63X oil immersion, NA=1.4, WD = 0.17 mm, Zeiss 44 07 61 1101-274) by selecting random fields of view containing multiple cells. Representative z-stacks were captured and 405, 488, 543, and 633 nm laser lines were used as appropriate for the immunofluorescence and nucleic acid stains. Image fields of view were collected until more than 50 cells were counted per condition. Human PMNs immunostained for PAD4 were imaged using a Nikon A1R confocal microscope equipped with a Nikon Apo 60x/1.4 oil λS DIC N2 objective lens and controlled by NIS-Elements.

##### Epifluorescence microscopy

dHL60 cells fixed after the NETosis endpoint assay (see above) were imaged using an Axiovert 200M wide-field fluorescence microscope (Zeiss) coupled to an AxioCam MR3 monochromatic CCD camera (Zeiss) using a Zeiss Plan-Neofluar 20x/0.4 Corr Ph2 objective lens with the Zeiss AxioVision software.

#### Image Analysis

##### Quantification of the percentage and timing of cellular events during NETosis

Time-lapse spinning disc confocal and DIC and movies of cells stained with SiR-DNA were used to quantify the percentage and timing of the following events: Plasma membrane microvesicle shedding, defined as the release of vesicles from the cell periphery as seen in DIC movies; DNA de-condensation, defined as the decrease in SiR-DNA inhomogeneity as seen in confocal fluorescence movies; Nuclear rounding, defined as the establishment of a circular nuclear periphery as seen in DIC and confocal fluorescence color overlay movies; DNA release from the nucleus, defined as expansion of the DNA periphery outside of the nuclear boundaries into the cytosol as seen in DIC and confocal fluorescence color overlay movies; PM permeabilization, defined as a decrease in contrast at the cell periphery as seen in DIC movies; and NETosis, defined as expansion of DNA outside of the cell boundary as seen in DIC and confocal fluorescence color overlay movies. All adherent cells in imaging fields were included in the analysis of the percentage of cells undergoing an event while only cells that remained in the field for the entire duration of the movie were included in the quantification of the timing of events. Multiple imaging fields were analyzed for each condition until the number of cells per condition was equal or greater than 50. The timing of an event was defined as the first time point when the corresponding event was observed in the movies by eye.

For quantification of the timing of cytoskeletal disassembly, vesiculation of the endoplasmic reticulum (ER), and rupture of the lamin meshwork, inner (Lap2β) and outer (ER) nuclear membranes in dHL60 cells, time-lapse spinning disc confocal DIC and fluorescence movies of cells transfected with the corresponding fluorescent-protein-tagged-cDNA were used. The timing of events was defined as follows; Actin disassembly: solubilization and decrease in the intensity of F-tractin-mApple; Microtubule disassembly: solubilization of Ensconsin-eGFP; Vimentin disassembly: solubilization of vimentin-mEmerald; ER vesiculation: appearance of discontinuities and vesicles in the ER network (stained with ER-5-mEmerald); Lamin A, B1, inner and outer nuclear membrane rupture: appearance of a discontinuity in the lamin meshwork or nuclear membrane. Cytoskeletal disassembly and ER vesiculation were detected using the confocal plane at the coverslip-cell interface while discontinuities in the lamin meshwork, inner and outer nuclear membranes were detected using the confocal plane at the cell center (3 µm above the cell-coverslip interface).

For quantification of the percentage and timing of cellular events in WT and PAD4 CR dHL60 cells transfected with mEmerald-C1, mEmerald-PAD4, mEmerald-PAD4-C645A or mEmerald-PAD4-K59A/K60A/K61A, only fluorescently labelled (and thus transfected) cells were taken into account.

##### Quantification of the Lamin meshwork rupture points

Time-lapse spinning disc confocal DIC and fluorescence movies of dHL60 cells transiently transfected with LaminA-mEmerald and stained with SiR-DNA were performed. Images were acquired every 1-2 min and the number of rupture points was defined as the number of discontinuities in the lamin A staining at the nuclear periphery the first time that the discontinuities were detectable by eye. Because the lamin meshwork in these cells is intrinsically discontinuous, only new discontinuities from where DNA expanded were taken into account.

##### Analysis of vimentin and lamin in mouse PMNs

Cells were counted from the fields in two independent experiments following an identical protocol with a DMSO vehicle control along with 5, 10, 60, and 240 minute time points. Images were evaluated for the distribution of vimentin immunofluorescence and the continuity of lamin immunofluorescence. The evaluations were done by two separate experimentalists.

##### Analysis of human neutrophils for PAD4 localization

For quantification of mean nuclear/cytosolic PAD4 ratio, the mean intensity of PAD4 in the nucleus (delineated by Hoechst DNA staining and DIC for nucleus outline) was divided by mean intensity of PAD4 in the cytosol (with reference to Hoechst DNA staining and DIC for cell outline) after each was corrected by background subtraction.

##### Quantification of plasma membrane permeabilization

Quantification of the ratio between the intracellular and extracellular calcein or dextran was performed by measuring the mean fluorescence intensity of the corresponding dye in a fixed-size ROI inside and outside the cell and calculating their ratio over time. Time of calcein and dextran entry was defined as the first time point (as detectable by eye) when the intracellular fluorescence of the corresponding dye increased. The DIC channel was used to define cells’ delimitations.

##### Quantification of Nuclear and cytosolic PAD4 and NLS

Measurement of the nuclear and cytoplasmic PAD4 and NLS intensity in dHL60 cells was performed using time-lapse movies of cells stained with SiR-DNA and transiently co-transfected with PAD4-mEmerald and NLS-mCherry. The confocal slice at the cell center (3 µm above the cell-coverslip interface) was used. Fixed-area ROIs inside the nucleus (as defined by the SIR-DNA staining) or in the cytosol (as defined by the DIC channel) were used to measure the mean PAD4 and NLS intensity overtime. The resulting mean intensity values were plotted after background subtraction and normalization to the maximum value of the corresponding mean intensity vector.

### Statistical analysis

All graphs were plotted using the GraphPad Prism software. Mann-Whitney and the Fisher’s Exact test were used to determine the significance of the difference between two distributions of random (timing graphs) or Boolean variables (percentages graphs). Mann-Whitney tests were performed using GraphPad Prism and Fisher’s Exact tests were done using R (version x64 3.5.1).

## Supplementary Figure Legends

**Figure S1:**
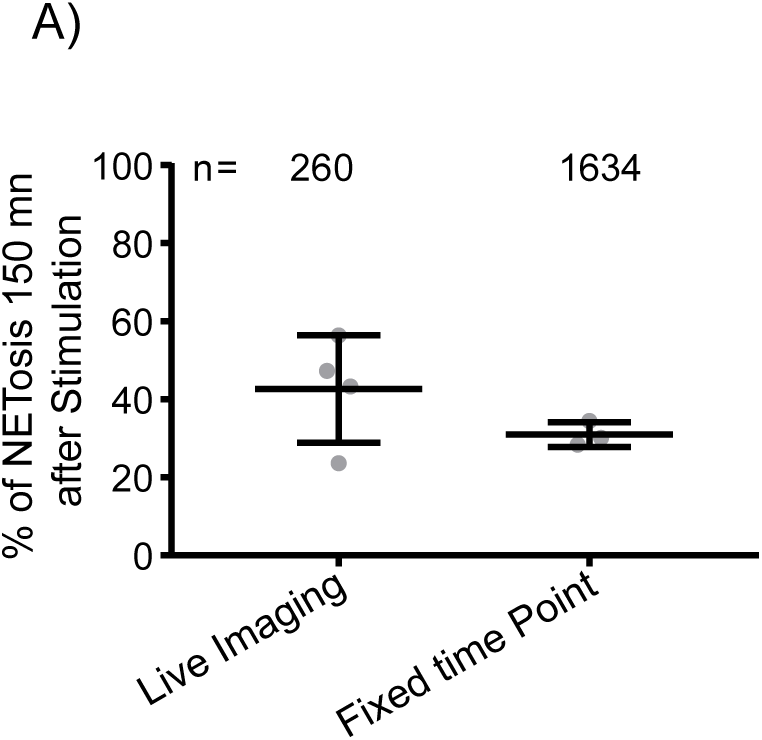
Live-cell imaging is minimally disruptive to NETosis. Differentiated HL60 neutrophil-like cells (dHL60) were plated on coverslips, stimulated with 4 μM ionomycin, fixed and stained with DAPI after 150 min or imaged live by differential interference contrast (DIC) and spinning-disk confocal microscopy at the coverslip-cell interface (Z = 0 μm) and 3 μm above in the cell center (Z = + 3 μm) at 1-2 min intervals for 4 hours. A: Quantification of the percentage of cells that released extracellular DNA after 150 min in immunostaining experiments (fixed time point) or live cell imaging. n = total number of cells observed, each point represents the percent of cells in one experiment. long bar = mean, short bars = S.D.

**Figure S2:**
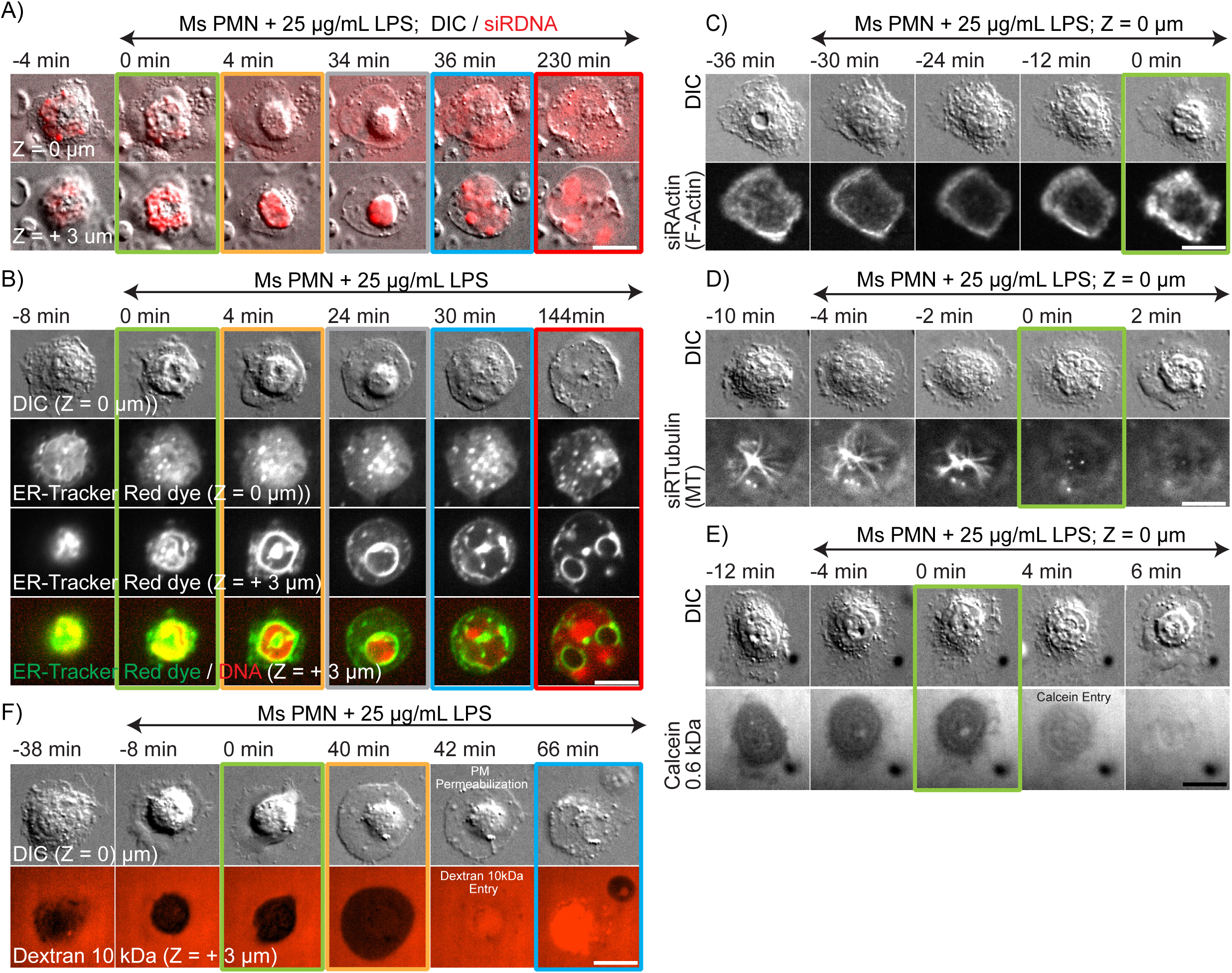
Cellular events mediating NETosis in mouse neutrophils stimulated with LPS. Mouse blood polymorphonuclear neutrophils (Ms PMN) were plated on coverslips, stimulated with 25 μg/mL of lipopolysaccharides (LPS), and imaged live by differential interference contrast (DIC) and spinning-disk confocal microscopy at the coverslip-cell interface (Z = 0 μm) and 3 μm above in the cell center (Z = + 3 μm) at 2 min intervals for 4 hours, with time in min relative to plasma membrane microvesicle shedding noted above images. Double-arrow-headed lines above images indicate cells that are in presence of ionomycin, boxes around images indicate different cellular events (green: MV Shedding; yellow: DNA de-condensation; grey: nuclear rounding; blue: DNA release to the cytosol, red: extracellular DNA release). A: Time series of image overlays of DIC (greyscale) and fluorescence (red) of cell and DNA dynamics in far-red SiR-DNA stained cells. B-F: Time-series of DIC (top row) and confocal (lower rows) images of live cells. B: Cells were stained with ER-Tracker Red (middle rows and bottom row, green) to label the ER and nuclear envelope (NE) and far red SiR-DNA (bottom rows, red) to label DNA. C, D: Cells were stained with far red SiR-actin to fluorescently label actin filaments (C) or SiR-tubulin to fluorescently label microtubules (D). E, F: soluble fluorescent markers (calcein (622 Da, E, greyscale) or Alexa fluor 594 10 kDa dextran (F, red)) were added to the imaging media to monitor changes in plasma membrane permeability. A-F: bars= 10 μm.

**Figure S3:**
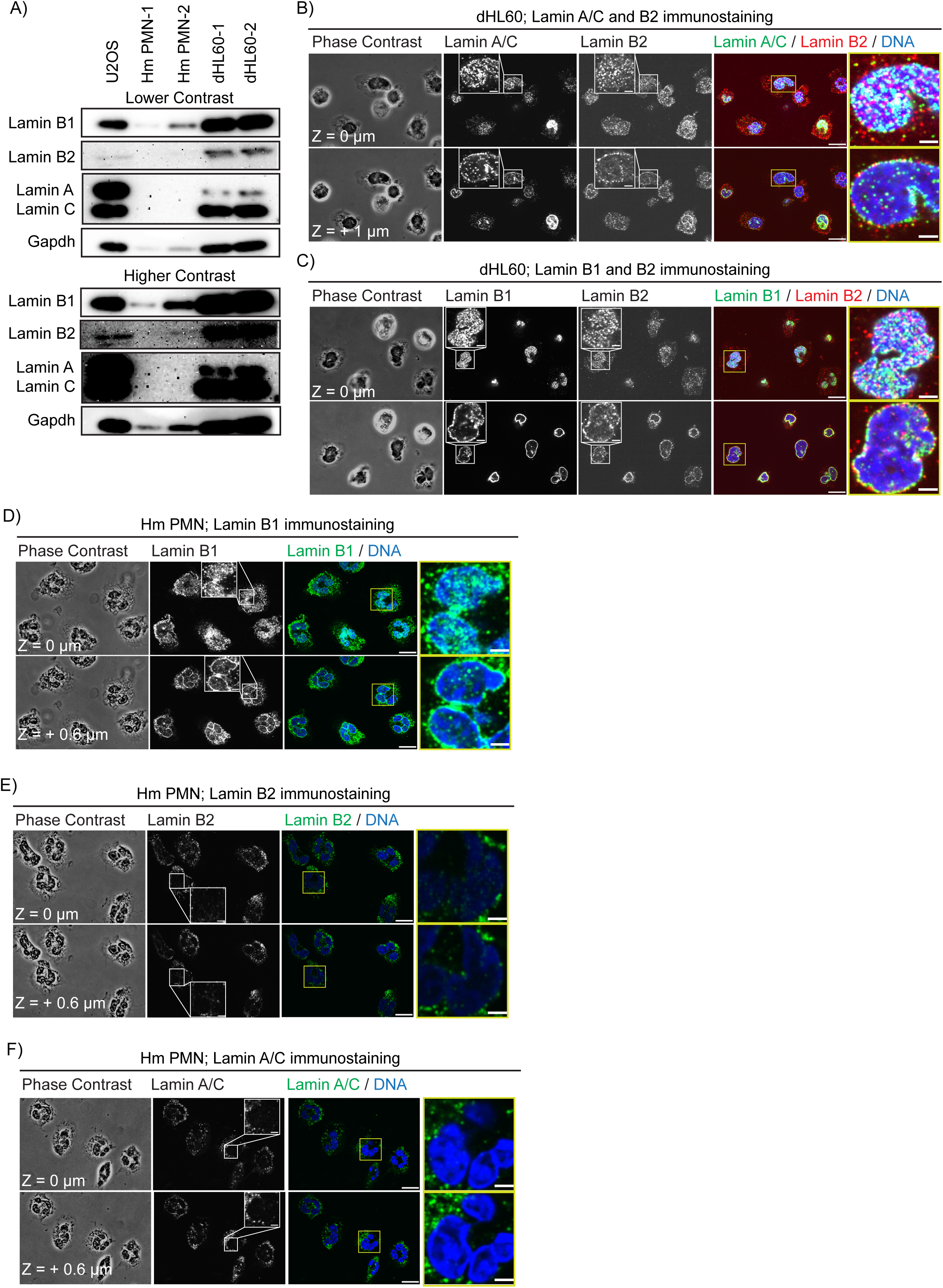
Expression and localization of lamins in human blood neutrophils and dHL60 cells. A: Western blot of lamin B1, B2 and A/C in lysates from human osteosarcoma cells (U2OS), human blood polymorphonuclear neutrophils from 2 different donors (Hm PMN-1 and Hm PMN-2) and differentiated HL60 neutrophil-like cells from 2 differentiation dates (dHL60-1 and dHL60-2). GAPDH was used as loading control. Two different contrast settings of the image are shown (upper panel, lower contrast, upper panel, higher contrast) to highlight bands of different abundance, B-F: Differentiated HL60 neutrophil-like cells (dHL60; B, C) or human blood polymorphonuclear neutrophils (Hm PMN; D, E, F) were plated on coverslips for 5 minutes then fixed and immunostained for lamin A/C, B1, B2 and DAPI to label DNA as noted on images. Z-stacks of 0.2 μm steps were acquired on a spinning-disk confocal microscope, and the Z distance from the coverslip surface is noted on the image. B-C: cells were co-immunostained with lamin B2 and A/C (B) or B1 (C). Insets show higher magnification of selected cells that are highlighted with a box. Color overlays are shown in the right-center and far right column, and the far right columns (outlined in yellow) show higher magnification of the boxed regions in the right-center columns. B-F: bars = 10 μm for lower magnification images and 2 μm for higher magnification images.

**Figure S4:**
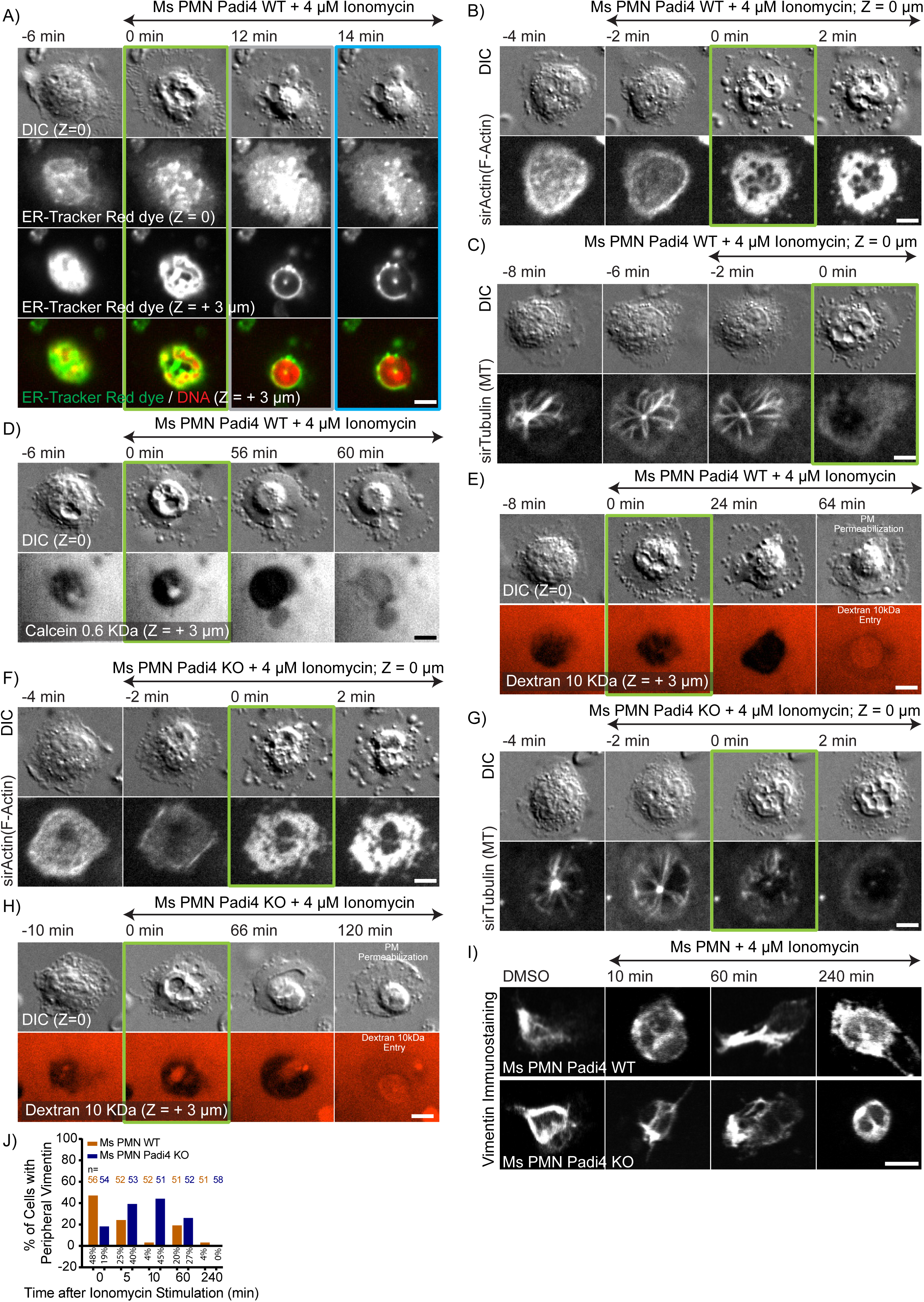
Cytoskeletal disassembly/remodeling, ER vesiculation and increase in plasma membrane permeability occur independent of PAD4 in mouse neutrophils. Mouse blood polymorphonuclear neutrophils (Ms PMN) from wild type (Padi4W T) or PAD4 knockout (Padi4 KO) were plated on coverslips, stimulated with 4 μM ionomycin, and fixed (I, J) at different time points after stimulation (noted on images, I) or imaged live (A-H) by differential interference contrast (DIC) and spinning-disk confocal microscopy at the coverslip-cell interface (Z = 0 μm) and 3 μm above in the cell center (Z = + 3 μm) at 2 min intervals for 4 hours, with time in min relative to plasma membrane microvesicle shedding noted on images. Double-arrow-headed lines above images indicate cells that are in presence of ionomycin, boxes around images indicate different cellular events (green: MV Shedding; blue: DNA release to the cytosol) Loss of DIC contrast indicating plasma membrane permeabilization and cellular entry of 10 kDa fluorescent dextran are noted on images in E and H. A-H: Time-series of DIC (top row) and confocal (lower rows) images of live cells. A: Cells were stained with ER-Tracker Red (middle two rows and bottom row, green) to label the ER and nuclear envelope (NE) and far red SiR-DNA (bottom rows, red) to label DNA. B, C, F, G: Cells were stained with far red SiR-actin to fluorescently label actin filaments (B, F) or SiR-tubulin to fluorescently label microtubules (C, G). D, E, H: soluble fluorescent markers (calcein (622 Da, D, greyscale) or Alexa fluor 594 10 kDa dextran (E, H, red)) were added to the imaging media to monitor changes in plasma membrane permeability. I: Cells were fixed at the times shown after ionomycin stimulation and vimentin was immunolocalized. J: Quantification from immunostaining experiments of cells with peripheral vimentin at different time points after ionomycin stimulation. A-I: bars= 5 μm.

**Figure S5:**
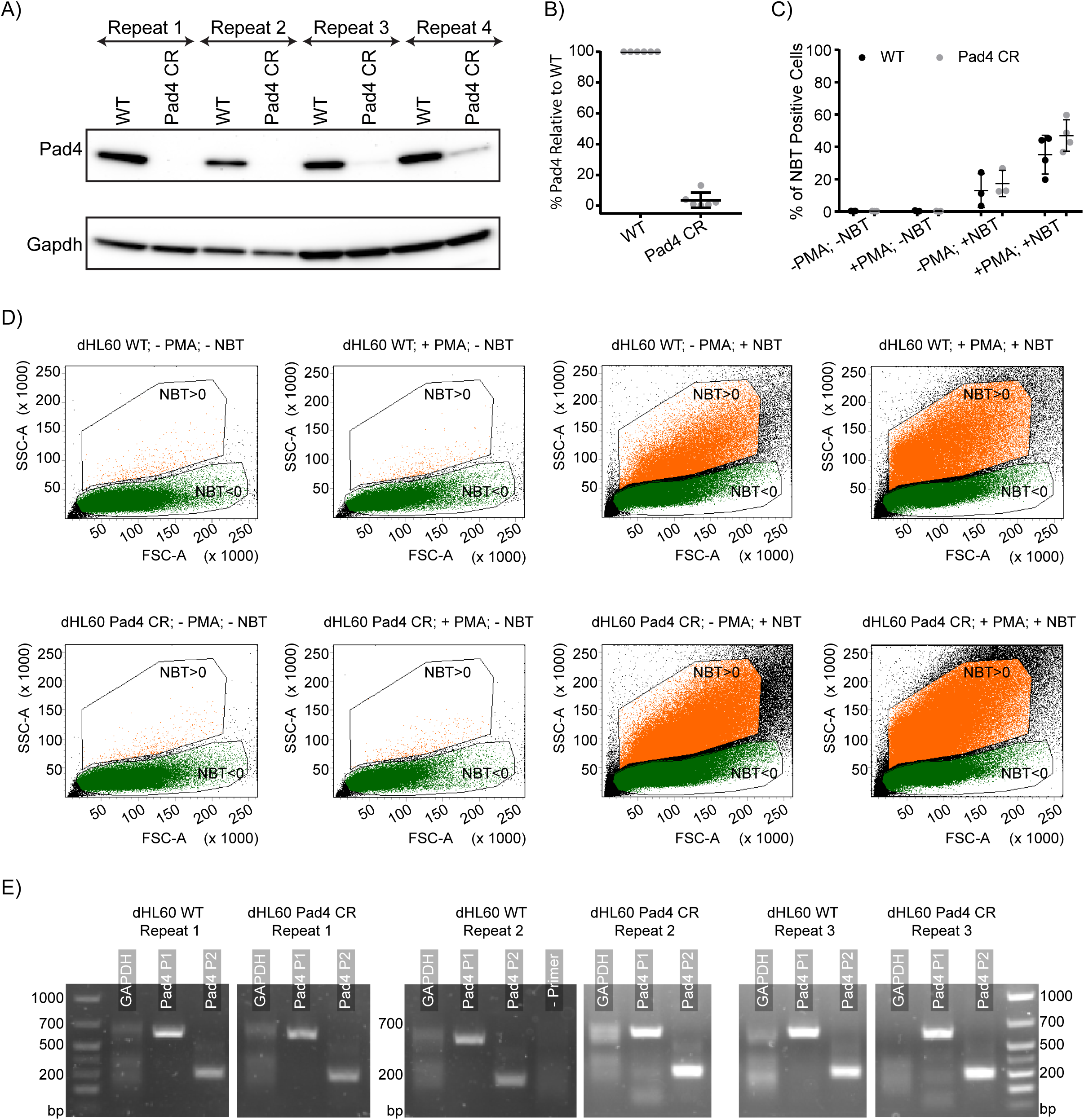
Characterization of the PAD4 CR dHL60 cell line. A: Western blot of PAD4 and GAPDH (as a loading control) for lysates of differentiated HL60 neutrophil-like cells (dHL60 WT) or dHL60 with the PAD4 gene modified by CRSPR-Cas9 gene editing (PAD4 CR dHL60) from four different differentiation dates (Repeat 1-4). B: Quantification from western blot experiments of the amount of PAD4 proteins expression relative to the WT levels in WT and PAD4 CR dHL60 cells, each point represents a distinct experiment. C, D: WT and PAD4 CR dHL60 cells were incubated in suspension for 25 min at 37C with DMSO or 20 nM Phorbol 12-myristate 13-acetate (PMA; +/-PMA) and +/-2 mg/mL of Nitro blue Tetrazolium Chloride (NBT; +/-NBT) then fixed before being analyzed by flow cytometry. C: Quantification of the percentage of NBT positive cells (cells in the NBT>0 gate in panel D) as determined by the increase in cells’ side-scatter signal. At least 100.000 cells were analyzed for each sample. D: Representative flow cytometry plots of cells’ side scatter area (SSC-A) as function of cells’ forward scatter area (FSC-A) that measure cell granularity and size respectively in WT (upper row) and PAD4 CR (bottom row) dHL60 cells. Green dots: cells that did not reduce NBT (NBT<0 gate); orange dots: cells that reduced NBT (NBT>0 gate; higher SSC-A value); black dots represent cells that are excluded from our gating strategy. From left to right: control cells, cells treated with PMA; cells incubated with NBT only; cells treated with PMA and incubated with NBT. E: RT-PCR of PAD4 and GAPDH mRNA in WT and PAD4 CR dHL60 cells from 3 distinct differentiation dates (Repeat 1-3) using 2 different sets of primers with expected product sizes of 610 (PAD4 P1) and 298 (PAD4 P2) bp, respectively.

## Supplementary Movie Legends

***Supplementary movie 1:*** Cell and DNA dynamics are similar in mouse and human PMN and dHL-60 cells during NETosis.

Mouse (Ms) or human (Hm) PMN or differentiated neutrophil-like (dHL-60) cells were stained with far-red SiR-DNA (red) and time-lapse DIC and spinning-disk confocal images were taken at one (dHL60) or two (Ms and Hm PMNs) min intervals at the coverslip surface (Z = 0 µm) and at three microns above (Z= +3 µm). Cells were stimulated with 4 µM ionomycin or 25 µg/mL of lipopolysaccharides (LPS) at the timepoint when the labels appear. Elapsed time shown in min relative to plasma membrane vesiculation. Scale bar = 5 μm.

***Supplementary movie 2:*** Cell, plasma membrane, and chromatin dynamics during NETosis in dHL-60 cells.

Differentiated neutrophil-like (dHL-60) cells were stained with far-red SiR-DNA (red, right panel) and co-transfected with the farnesylation signal sequence CAAX fused to mApple (CAAX-mApple, center panel) as a plasma membrane marker and histone H1 fused to mEmerald (H1-mEm, green right panel) and time-lapse DIC and spinning-disk confocal images were taken every min. Cells were stimulated with 4 µM ionomycin at the timepoint when the label appears. Elapsed time shown in min relative to plasma membrane vesiculation. Scale bar = 10 μm.

***Supplementary movie 3:*** Rapid drop in cytoplasmic refractive index suggests plasma membrane permeabilization during NETosis dHL-60 cells.

Differentiated neutrophil-like (dHL-60) cells were stained with far-red SiR-DNA (red) and time-lapse DIC and spinning-disk confocal images were taken every min at three microns above the coverslip surface (Z= +3 um). Cells were stimulated with 4 µM ionomycin at the timepoint when the label appears. Right panel shows a zoom of the image on the left. Timepoint of drop in refractive index noted (PM Permeabilization). Elapsed time shown in min relative to plasma membrane vesiculation. Scale bar = 5 μm (left) or 2 µm (right).

***Supplementary movie 4:*** Lamin meshwork rupture during NETosis in dHL-60 cells.

Differentiated neutrophil-like (dHL-60) cells were stained with far-red SiR-DNA (red, right panels) and transfected with either mEmerald (green) fused to lamin A (upper row) or lamin b (lower row). Time-lapse DIC and spinning-disk confocal images were taken at one min (for lamin A) or two min (for lamin B1) min intervals at three microns above the coverslip surface (Z= +3 µm). Cells were stimulated with 4 µM ionomycin at the timepoint when the label appears. Elapsed time shown in min relative to plasma membrane vesiculation. Scale bar = 5 μm.

***Supplementary movie 5:*** Endoplasmic reticulum (ER) vesiculation during NETosis in mouse and human PMN and dHL-60 cells.

Mouse (Ms) or human (Hm) PMN were stained with ER-Tracker Red vital dye and differentiated neutrophil-like (dHL-60) cells were transfected with the KDEL sequence combined with the ER-retention signal sequence from calreticulin fused to mEmerald (ER-5-mEmerald) and time-lapse DIC and spinning-disk confocal images were taken at 2 min (for human and mouse PMN) or 1 min (for dHL-60) intervals at the coverslip surface (Z = 0 µm). Cells were stimulated with 4 µM ionomycin or 25 µg/mL lipopolysaccharides (LPS) at the timepoint when the labels appear. Elapsed time shown in min relative to plasma membrane vesiculation. Scale bars = 10 μm.

***Supplementary movie 6:*** Nuclear envelope rupture during NETosis in human PMN and dHL-60 cells.

Human (Hm) PMN were stained with far-red SiR-DNA (Red, right column) and ER-Tracker Red vital dye (green bottom left) and differentiated neutrophil-like (dHL-60) cells were stained with far-red SiR-DNA (Red, right column) and transfected with either the inner nuclear envelope protein Lap2β fused to mEmerald (green, upper right) or the KDEL sequence combined with the ER-retention signal sequence from calreticulin fused to mEmerald (ER-mEmerald, green, middle right). Time-lapse DIC and spinning-disk confocal images were taken at one min (for Lap2β-mEmerald and ER-mEmerald) or two min (for ER-tracker dye) at 3 µm above the coverslip surface (Z = +3 um). Cells were stimulated with 4 uM ionomycin at the timepoint when the labels appear. Arrowheads highlight rupture initiation points. Elapsed time shown in min relative to plasma membrane vesiculation. Scale bars = 5 µm (top and middle row) or 10 μm (bottom row).

***Supplementary movie 7:*** Nuclear envelope and lamin meshwork rupture during NETosis in dHL-60 cells.

Differentiated neutrophil-like (dHL-60) cells stained with far-red SiR-DNA (blue) and co-transfected with the inner nuclear envelope protein Lap2β fused to mEmerald (green) and lamin B1 fused to mApple (red) and time-lapse DIC and spinning-disk confocal images were taken every min at 3 µm above the coverslip surface (Z = +3 µm). Cells were stimulated with 4 µM ionomycin at the timepoint when the labels appear. Arrowheads highlight rupture initiation points. Elapsed time shown in min relative to plasma membrane vesiculation. Scale bar = 5 μm.

***Supplementary movie 8:*** Cytoskeletal remodeling and disassembly during NETosis in dHL-60 cells.

Differentiated neutrophil-like (dHL-60) cells were transfected with F-tractin-mApple to mark actin filaments, vimentin-mEmerald, or the microtubule-binding domain of ensconsin fused to eGFP (Ensconsin-eGFP) to mark microtubules. Time-lapse DIC and spinning-disk confocal images were taken at 15 sec (for F-tractin) or one min (for vimentin and ensconsin) intervals at the coverslip surface (Z = 0 µm). Cells were stimulated with 4 µM ionomycin at the timepoint when the labels appear. Elapsed time shown in min relative to plasma membrane vesiculation. Scale bar = 5 μm.

***Supplementary movie 9:*** Cytoskeletal remodeling and disassembly during NETosis in human PMN.

Human (Hm) PMN were stained with far-red SiR-Actin or SiR-Tubulin to label actin and microtubules, respectively, and time-lapse DIC and spinning-disk confocal images were taken every 2 min at the coverslip surface (Z = 0 µm). Cells were stimulated with 4 µM ionomycin at the timepoint when the labels appear. Elapsed time shown in min relative to plasma membrane vesiculation. Scale bar = 10 μm.

***Supplementary movie 10:*** Cytoskeletal remodeling and disassembly during NETosis in mouse PMN.

Mouse (Ms) PMN were stained with far-red SiR-Actin or SiR-Tubulin to label actin and microtubules, respectively, and time-lapse DIC and spinning-disk confocal images were taken every 2 min at the coverslip surface (Z = 0 µm). Cells were stimulated with 4 µM ionomycin or 25 µg/ml lipopolysaccharides (LPS) at the timepoint when the labels appear. Elapsed time shown in min relative to plasma membrane vesiculation. Scale bar = 10 μm.

***Supplementary movie 11:*** Jasplakinilode blocks actin disassembly and inhibits NET release in dHL-60 cells.

Differentiated neutrophil-like (dHL-60) cells were transfected with actin-mEmerald (G-actin-mEmerald, green) and stained with far-red SiR-DNA (red) and time-lapse DIC and spinning-disk confocal images were taken every two min at the coverslip surface (Z = 0 µm) and three microns above the coverslip surface (Z = +3 µm) in the absence (top row) or presence (bottom two rows) of 1 µM jasplakinolide. Two examples of jasplakinolide-treated cells are shown: one in which the cortical actin blocks NET release (middle row) and one in which the NET is released at a gap in the cortical actin (bottom row). Cells were stimulated with 4 µM ionomycin at the timepoint when the labels appear. Elapsed time shown in min relative to plasma membrane vesiculation. Scale bar = 5 μm.

***Supplementary movie 12:*** Taxol blocks microtubule disassembly but does not affect NET release in dHL-60 cells.

Differentiated neutrophil-like (dHL-60) cells were transfected with tubulin-mEmerald (green) and stained with far-red SiR-DNA (red) and time-lapse DIC and spinning-disk confocal images were taken every two min at the coverslip surface (Z = 0 µm) and three microns above the coverslip surface (Z = +3 µm) in the absence (top row) or presence (bottom row) of 10 µM taxol. Cells were stimulated with 4 µM ionomycin at the timepoint when the labels appear. Elapsed time shown in min relative to plasma membrane vesiculation. Scale bar = 5 μm.

***Supplementary movie 13:*** Plasma membrane ruptures at NET release in dHL-60 cells.

Differentiated neutrophil-like (dHL-60) cells were transfected with the farnesylation signal sequence CAAX fused to mApple (CAAX-mApple) as a plasma membrane marker and stained with far-red SiR-DNA (red) and time-lapse DIC and spinning-disk confocal images were taken every two min at three microns above the coverslip surface (Z = +3 µm). Cells were stimulated with 4 µM ionomycin at the timepoint when the labels appear. Elapsed time shown in min relative to plasma membrane vesiculation. Scale bar = 5 μm.

***Supplementary movie 14:*** Plasma membrane permeabilizes prior to NET release in dHL-60 cells.

Differentiated neutrophil-like (dHL-60) cells were subjected to time-lapse DIC and spinning-disk confocal images taken every two min at three microns above the coverslip surface (Z = +3) in media containing calcein (622 Da, top row) or Alexa fluor 647 10 kDa dextran (bottom row). Cells were stimulated with 4 µM ionomycin at the timepoint when the labels appear. Elapsed time shown in min relative to plasma membrane vesiculation. Scale bar = 5 μm.

***Supplementary movie 15:*** Stepwise increase in plasma membrane permeability prior to NET release in dHL-60 cells.

Differentiated neutrophil-like (dHL-60) cells were subjected to time-lapse DIC and spinning-disk confocal images taken every one or two min at the the coverslip surface (Z = 0 µm) in media containing calcein (622 Da) and Alexa fluor 594 10 kDa dextran (red, top row) or Alexa fluor 647 10 kDa dextran (red) and Oregon Green 488 70 kDa (green, bottom row). Cells were stimulated with 4 µM ionomycin at the timepoint when the labels appear. Elapsed time shown in min relative to plasma membrane vesiculation. Scale bar = 10 μm.

***Supplementary movie 16:*** Comparison of cell and DNA dynamics in wild type and PAD4 KO mouse PMN during NETosis.

Wild-type (WT) and PAD4 knockout (Padi4 KO) mouse (Ms) PMN were stained with far-red SiR-DNA (red) and time-lapse DIC and spinning-disk confocal images were taken at two min intervals at the coverslip surface (Z = 0 µm) and at three microns above (Z= +3 µm). Cells were stimulated with 4 µM ionomycin at the timepoint when the labels appear. Elapsed time shown in min relative to plasma membrane vesiculation. Scale bar = 5 μm.

***Supplementary movie 17:*** Comparison of endoplasmic reticulum and nuclear envelope dynamics in wild type and PAD4 KO mouse PMN during NETosis.

Wild-type (WT) and PAD4 knockout (Padi4 KO) mouse (Ms) PMN were stained with far-red SiR-DNA (red) and ER-Tracker Red vital dye (green) and time-lapse DIC and spinning-disk confocal images were taken at two min intervals at the coverslip surface (Z = 0 µm) and at three microns above (Z= +3 µm). Cells were stimulated with 4 µM ionomycin at the timepoint when the labels appear. Elapsed time shown in min relative to plasma membrane vesiculation. Scale bar = 5 μm.

***Supplementary movie 18:*** PAD4 dynamics in dHL-60 cells during NETosis.

Differentiated neutrophil-like (dHL-60) cells were transfected with PAD4-mEmerald (green) and mCherry fused to the nuclear localization signal sequence which served as a soluble nucleoplasmic marker (NLS-mCherry) and stained with far-red SiR-DNA (blue). Time-lapse DIC and spinning-disk confocal images were taken every two min at the coverslip surface (Z = 0 µm) and at three microns above the coverslip surface (Z = +3 µm). Cells were stimulated with 4 µM ionomycin at the timepoint when the labels appear. Elapsed time shown in min relative to plasma membrane vesiculation. Scale bar = 10 μm.

***Supplementary movie 19:*** Comparison of cell and DNA dynamics in wild type and PAD4 CR dHL-60 cells during NETosis.

Differentiated neutrophil-like (dHL-60) cells (WT) and dHL-60 cells with PAD4 modified by gene editing (Pad4 CR) were stained with far-red SiR-DNA (red) and time-lapse DIC and spinning-disk confocal images were taken at one min intervals at the coverslip surface (Z = 0 µm) and at three microns above (Z= +3 µm). Cells were stimulated with 4 µM ionomycin at the timepoint when the labels appear. Elapsed time shown in min relative to plasma membrane vesiculation. Scale bar = 10 μm.

***Supplementary movie 20:*** The enzymatic and nuclear localization activities of PAD4 are required for NETosis in dHL-60 cells.

Differentiated neutrophil-like cells with PAD4 modified by gene editing (dHL60 Pad4 CR) were transfected with mEmerald (mEmerald-C1, green, first column) or mEmerald-tagged PAD4 (mEmerald Pad4, green, second column) or mEmerald-tagged PAD4 variants bearing point mutations coding amino acid substitutions that specifically inactivate its enzymatic activity (mEmerald-PAD4-C645A, green, third column) or nuclear localization signal (mEmerald-Pad4-K59A/K60A/K61A) were stained with far-red SiR-DNA (red) and time-lapse DIC and spinning-disk confocal images were taken at two min intervals at the coverslip surface (Z = 0 µm) and at three microns above (Z= +3 µm). Cells were stimulated with 4 µM ionomycin at the timepoint when the labels appear. Elapsed time shown in min relative to plasma membrane vesiculation. Scale bar = 10 μm.

